# *RAS*-mutant clones drive extramedullary acute myeloid leukemia

**DOI:** 10.64898/2026.04.07.715220

**Authors:** Panagiota Chaida, Julia Frimmel, Lara Hopfer, Bianca Perfler, Eva Gruden, Akshaya Kailasnathan, Karin Lind, Bernadette Bramreiter, Johannes Foßelteder, Sonja Wurm, Jennifer Neiss, Stefan Köck, Dominik Wolf, Gudrun Ratzinger, Nassim Ghaffari-Tabrizi-Wizsy, Beate Rinner, Karoline Fechter, Kristina Glebova, Gudrun Pregartner, Katarina Vizar Cisarova, Gerald Hoefler, Karl Kashofer, Andreas Prokesch, Annkristin Heine, Albert Wölfler, Heinz Sill, Andreas Reinisch, Friedrich Stölzel, Armin Zebisch

## Abstract

Extramedullary acute myeloid leukemia (eAML) represents a clinically challenging manifestation of acute myeloid leukemia (AML), but its molecular drivers remain poorly defined. We performed targeted sequencing in 85 eAML biopsies, representing one of the largest molecular analyses of eAML to date. We detected mutations in *RAS* or *RAS*-modifying genes (*RAS*^*MUT*^; *NRAS, KRAS, PTPN11, CBL*, and *NF1*) in 41% of cases, representing a significant enrichment compared to bone marrow (BM) samples of more than 1300 AML patients not selected for eAML. Analysis of paired eAML and BM specimens revealed expansion and/or de-novo appearance of *RAS*^*MUT*^ clones at the extramedullary site. Functional studies using primary murine leukemia cells and CRISPR/Cas9-engineered isogenic human leukemia cell lines demonstrated that *RAS*^*MUT*^ increase the migration and invasion of leukemic cells compared to *RAS*-wildtype controls. Consistently, *RAS*^*MUT*^ cells showed increased infiltration into the chorioallantoic membrane of chicken embryos and demonstrated enhanced extramedullary growth after injection into immunocompromised mice. RNA sequencing revealed increased expression of junctional adhesion molecule-like (*JAML*) and activation of PI3K/AKT signaling in *RAS*^*MUT*^ cells. *JAML* silencing and pharmacologic AKT inhibition reversed the *RAS*^*MUT*^-driven effects on leukemic cell migration, demonstrating a causal role of the JAML-PI3K/AKT axis in *RAS*^*MUT*^-driven eAML formation. In conclusion, these findings delineate the molecular landscape of extramedullary AML and show that *RAS*^*MUT*^ are enriched within this AML subform. They further demonstrate that *RAS*^*MUT*^ actively contribute to leukemic tissue infiltration through activation of a *RAS*^*MUT*^-JAML-PI3K/AKT axis, highlighting AKT signaling as a potential therapeutic vulnerability in *RAS*^*MUT*^-associated eAML.

## Introduction

Extramedullary acute myeloid leukemia (eAML) is a specific manifestation of acute myeloid leukemia (AML), where myeloid blasts invade extrahematopoietic tissues. Approximately 9% of AML cases exhibit additional eAML,^1^ although newer studies incorporating PET scans suggest significantly higher numbers.^2,3^ Data about the clinical relevance of eAML have mainly been derived from retrospective and observational trials and have remained controversial for a long time^4,5^. However, newer meta-analyses and studies including patients’ genetic profile suggest that the prognostic relevance may depend on the affected eAML site and its molecular profile.^3,4,6,7^ They also show that eAML may be insufficiently cleared by currently used therapeutic regimens and serve as a leukemic reservoir that seeds systemic relapse.^8–10^ Despite this evidence, most knowledge about the molecular profile of eAML is derived from corresponding BM specimens, and sequencing studies from eAML biopsies are rare and restricted to small cohort sizes. Consequently, the knowledge about the molecular pathogenesis and relevant eAML disease drivers is insufficient.

In this study, we performed targeted sequencing in 85 eAML biopsies and identified mutations in *RAS* or *RAS*-modifying genes (*RAS*^*MUT*^; *NRAS, KRAS, CBL, PTPN11, NF1*) in 41% of cases. *RAS*^*MUT*^ were not only enriched in eAML but also showed clonal expansion at the extramedullary sites. Furthermore, we show that *RAS*^*MUT*^ are functionally involved in the tissue infiltration of leukemic blasts and eAML formation. Finally, we elaborate on the mechanisms and show that junctional adhesion molecule-like protein (JAML) mediated PI3K/AKT activation is a central step in *RAS*^*MUT*^-mediated extramedullary leukemia growth and represents an interesting therapeutic target for *RAS*^*MUT*^ eAML manifestations.

## Materials and methods

### Patient samples, murine models, and cell lines

Formalin-fixed paraffin-embedded (FFPE) eAML patient specimens and BM aspirates were collected at the Medical University of Graz (MUG, provided by the Biobank of the MUG, Cohort 6002_14, Myeloid Neoplasias Collection),^11–13^ the Medical University of Innsbruck, the University Hospitals Schleswig-Holstein-Kiel and Dresden (both in Germany) between 2003 and 2025. Additional AML BM controls (AML, LB-MUG cohort, Figure 1C) were collected at the MUG and partly published previously.^14^ The study was approved by the local institutional review boards (Graz: 1212/2024 and 30464ex17/18; Innsbruck: 1217/2025; Germany-TU Dresden EK98032010) and performed in accordance with the Declaration of Helsinki. Murine hematopoietic stem and progenitor cells (HSPC) for in vitro assays were collected from 8-9 week-old male and female Mx1-Cre-*Kras*^*G12D/Wt*^ and Mx1-Cre-*Kras*^*Wt/Wt*^ control mice on a C57BL/6 strain background as described.^15,16^ NOD-Rag2tm1-Il2rgtm1-Foxn1nu/Rj (NRG) nude mice were obtained from Janvier Labs (Le Genest-Saint-Isle, France). All mouse experiments were approved by the Federal Ministry of Education, Science and Research in Austria (GZ: 20240691.181 and BMWFW-66.010/0046-WF/V/3b/2016). Cell lines were obtained from the German National Resource Center for Biological Material (DSMZ, Braunschweig, Germany).

**Figure 1.**
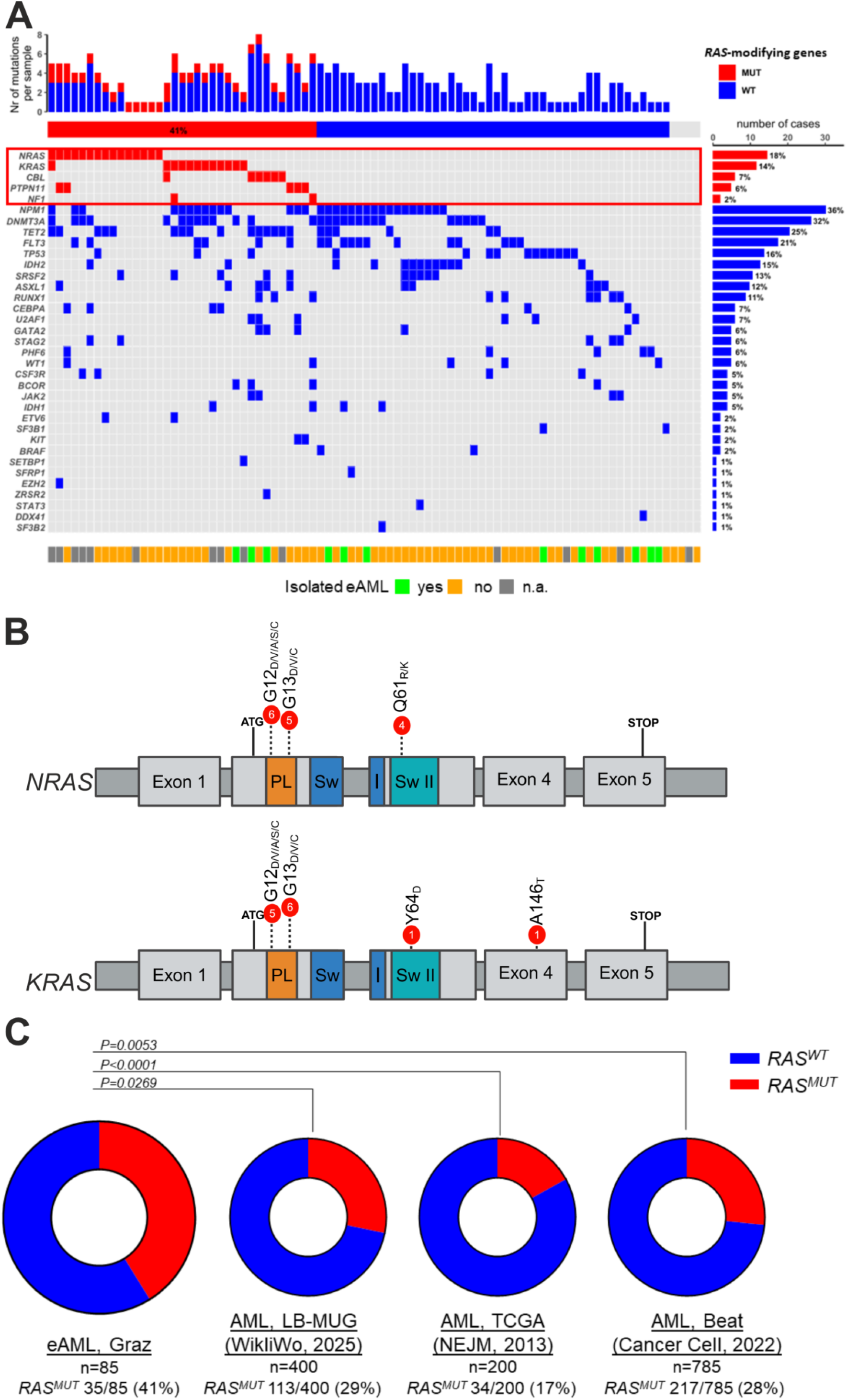
*RAS* and *RAS*-modifying mutations *(RAS*^*MUT*^) are enriched in extramedullary acute myeloid leukemia (eAML). (A) Oncoplot illustrating the molecular landscape of 85 eAML patients. Each column on the x-axis represents an individual patient. *RAS*^*MUT*^ are shown in red, *RAS*-wildtype (*RAS*^*WT*^) cases in blue. Only genes harboring at least one mutation are displayed. (B) Lollipop plot showing hotspot mutation codons in *NRAS* and *KRAS*. The height of each bar and the number within the red circle represent the number of occurrences of the respective mutation. (C) Donut plots representing the frequency of *RAS*^*MUT*^ (red) and *RAS*^*WT*^ (blue) in eAML compared with three independent acute myeloid leukemia (AML) bone marrow cohorts (LB-MUG^14^, TCGA^25^ and Beat-AML^24^). Statistical differences were calculated using the Fisher’s exact test. Nr, number; MUT, mutant; WT, wildtype; n.a., not available; PL, P-loop protein domain; SwI, Switch I protein domain; SwII, Switch II protein domain; TCGA, The Cancer Genome Atlas; LB-MUG, Leukemia Biobank-Medical University of Graz.

### Targeted next generation sequencing (NGS)

For Austrian samples, eAML in FFPE specimens was initially confirmed by histology and immunohistochemistry. Subsequently, NGS was performed as previously described, covering a set of 49 genes recurrently mutated in myeloid neoplasms (Supplemental methods).^14,17^ For the German specimens, snap-frozen tissue samples with immunohistochemistry-confirmed eAML were analyzed using capture-based targeted NGS with the Myeloid-NGS DNA Capture (Myeloid-NDC, Univ8 Genomics, Belfast, United Kingdom), and sequence variants were annotated.

### CRISPR/Cas9 mutational knock-in and siRNA transfection

K562 and HEL hematopoietic cell lines were maintained at 37°C/5% CO_2_ in RPMI-1640 enriched with 10% fetal bovine serum (FBS) and 100U/mL penicillin, 100mg/mL streptomycin and 0.25mg/mL amphotericin B (Thermo Fisher Scientific, Waltham, MA, USA). *NRAS*^*G12D/WT*^ and *NRAS*^*WT/WT*^ cell lines were generated using CRISPR-Cas9-mediated knock-in as previously described^18,19^ and as detailed in the Supplemental methods. Transient knockdown of *JAML* was performed in K562-*NRAS*^*G12D/WT*^ cells with two different ON-TARGETplus siRNA sequences and a non-targeting control (Dharmacon, Lafayette, CO; Revvity, Waltham, MA, USA) as previously described^11,20^ and as detailed in the Supplemental methods.

### Migration and invasion assays

Transwell migration and invasion assays were performed as previously described^21^ using Corning Transwell migration (3µm pores) and Matrigel-coated invasion chambers (8µm pores; both Corning Inc., Corning, NY, USA). A total of 1.5×10^5^ cells in suspension in serum-free RPMI-1640 medium was added to the upper chamber inserts, which were placed into wells containing complete medium supplemented with 10% FBS as a chemoattractant. Migration/invasion was assessed after 24h (or after 12h in experiments following siRNA transfection).

### Chorioallantoic membrane (CAM) assays

Ex ovo CAM assays in chicken embryos were performed as described previously.^21^ Briefly, K562-*NRAS*^*G12D/WT*^ and K562-*NRAS*^*WT/WT*^ cells (1×10^6^ cells in 15μl PBS) mixed with 5μl Matrigel ® (Corning® Basement Membrane Matrix) were placed directly onto the CAM at day 10 of embryonic development. Four days later, onplant regions of the CAM were harvested and analyzed for cell infiltration and tumor formation. Tissue sections (5µm) were prepared, and sections containing thickest tumor areas were stained with hematoxylin and eosin (H&E). Fluorescence signals were analyzed using ImageJ (https://imagej.net/ij/)

### NRG nude mouse assays

5×10^6^ cells of K562-*NRAS*^*G12D/WT*^ and K562-*NRAS*^*WT/WT*^ were injected subcutaneously into male and female 6-week-old NRG nude mice into both flank regions of the same animal under anesthetized conditions (n=7-10). Tumor growth (length, width, and height) was measured twice weekly via ultrasound (US) (tumor volume in mm^3^=[(length)x(width)x(height)]x(π/6)]. Due to rapid tumor growth in the *NRAS*^*G12D/WT*^ group already at day 14 after injection, mice were sacrificed in accordance with ethical regulations. Tumors were excised, weighed, and shock frozen in methylbutan, followed by storage in liquid nitrogen until RNA-Seq analysis. Peripheral blood and BM aspirates were analyzed by flow cytometry to detect circulating tumor cells based on fluorescent protein expression.

### Whole transcriptome RNA sequencing (RNA-Seq)

RNA extraction from nude mice tumors and RNA-Seq was conducted as a service by Azenta Life Sciences (Griesheim, Germany), and raw reads in FASTQ format were provided. Data analysis and gene set enrichment analysis are described in Supplemental Methods. All raw data are publicly available via the NCBI Gene Expression Omnibus (accession number will be provided upon publication).

### Database analysis and statistical analysis

Sequencing data for *RAS*^*MUT*^ status were obtained from cBioportal^22,23^ (v6.0.27) for Cancer Genomics from the following datasets: Beat-AML^24^ (downloaded 20-NOV-2025) and TCGA-LAML^25^ (downloaded 20-NOV-2025). RNA-Seq data for gene expression analyses within the Beat-AML cohort were downloaded 16-JAN-2025 using mRNA expression z-scores relative to all samples (log RNA-Seq RPKM). For correlation analyses between *JAML* expression, leukocyte migration and PI3K/AKT pathway activity, gene set–based signature scores were calculated using curated gene sets from the Molecular Signatures Database^26,27^ (MSigDB, v2026.1) on 13-FEB-2026. The PI3K/AKT activation score was derived from the HALLMARK_PI3K_AKT_MTOR_SIGNALING gene set.^28^ Leukocyte migration scores were calculated based on genes annotated to the Gene Ontology term GO:0050900 (leukocyte migration).^26^ All Statistical analyses were performed using R, version 4.5.0; https://www.R-project.org/ and GraphPad Prism version 10.5.0 (San Diego, CA, USA). All p-values are two-sided; the significance level was 0.05. The specific statistical tests used are indicated in the respective figure legends.

## Results

### *RAS*^*MUT*^ are frequently detected in eAML and enriched at the extramedullary sites

In order to shed more light on the molecular pathogenesis of eAML we performed targeted sequencing of 49 myeloid neoplasm-associated genes in eAML biopsies of 85 patients. Patients were diagnosed at four Austrian and German hematology centers (Graz, n=53; Innsbruck, n=7; Dresden, n=22; and Schleswig-Holstein-Kiel, n=3. Of those with available data (n=71), twelve had isolated eAML and 59 eAML coinciding with leukemic BM infiltration. Detailed clinical characteristics and eAML localizations are depicted in Supplemental Table 1. Mutation calling necessitated a variant allele frequency (VAF) of at least 5% with a 1000X coverage. However, lower VAF cutoffs were accepted when the same mutation was detected in the corresponding BM specimen.^29^ Our analyses validated the high frequency of mutations in *NPM1* (26%), *DNMT3A* (21%), *TET2* (19%), and *FLT3* (16%).^30,31^ In addition, 41% of patients exhibited one or more *RAS*^*MUT*^, most frequently affecting *NRAS* (18%), followed by *KRAS* (14%), *CBL* (7%), *PTPN11* (6%) and *NF1* (2%; Figure 1A). The most frequently affected region in *NRAS* was codon 12 (40%), followed by variants affecting codon 13 (33%) and 61 (27%; Figure 1B). The most frequently affected regions in *KRAS* were codon 13 (46%) and 12 (38%; Figure 1B). We then compared the frequency of *RAS*^*MUT*^ in this eAML cohort with *RAS*^*MUT*^ frequencies in BM specimens from three cohorts of patients not selected for eAML (n=1.385; including a local cohort from the Medical University of Graz [LB-MUG],^14^ n=400; TCGA-LAML,^25^ n=200; Beat-AML,^24^ n=785). *RAS*^*MUT*^ were enriched in the eAML specimens compared to all three cohorts studied (Figure 1C). Even when the three groups were combined, *RAS*^*MUT*^ were enriched in eAML (*RAS*^*MUT*^ 33/85 [41%] in eAML vs 364/1385 [26%] in the AML patients not selected for eAML; P=0.016).

Next, we assessed paired NGS results of *RAS*^*MUT*^ eAML biopsies and affected BM of the corresponding patients. These analyses were possible in 25 *RAS*^*MUT*^ patients. Out of these, four patients developed a *RAS*^*MUT*^ in the eAML specimen that was not present in the BM. We aimed to compare the VAFs between BM and eAML specimens. Therefore, we employed a previously developed normalization strategy to account for differences in leukemia cell burden across BM and eAML tissues.^32^ In brief, each mutation’s VAF within a sample was normalized relative to the highest VAF mutation in that sample, enabling a standardized assessment of clonal proportions irrespective of absolute VAF numbers. These normalized *RAS*^*MUT*^ VAFs in the eAML specimens were significantly higher than those in the corresponding AML BM (70% vs 16%, P=0.003; Figure 2A). When looking at individual samples, normalized *RAS*^*MUT*^ VAF increased in the eAML specimen in 18/25 (72%) of cases, remained unchanged in 5/25 (20%) cases, and decreased in only 2/25 (8%; Figure 2B). Consistently, not a single *RAS*^*WT*^ eAML case with matched BM available for analysis (n=40) exhibited a loss of the *RAS* mutation at the extramedullary site (Supplemental Table 2). Representative examples of clonal evolution during eAML development are shown in Figure 2C, illustrating the progressive expansion of the *RAS*-mutant population and a corresponding decline in the *RAS*-wildtype clones.

**Figure 2.**
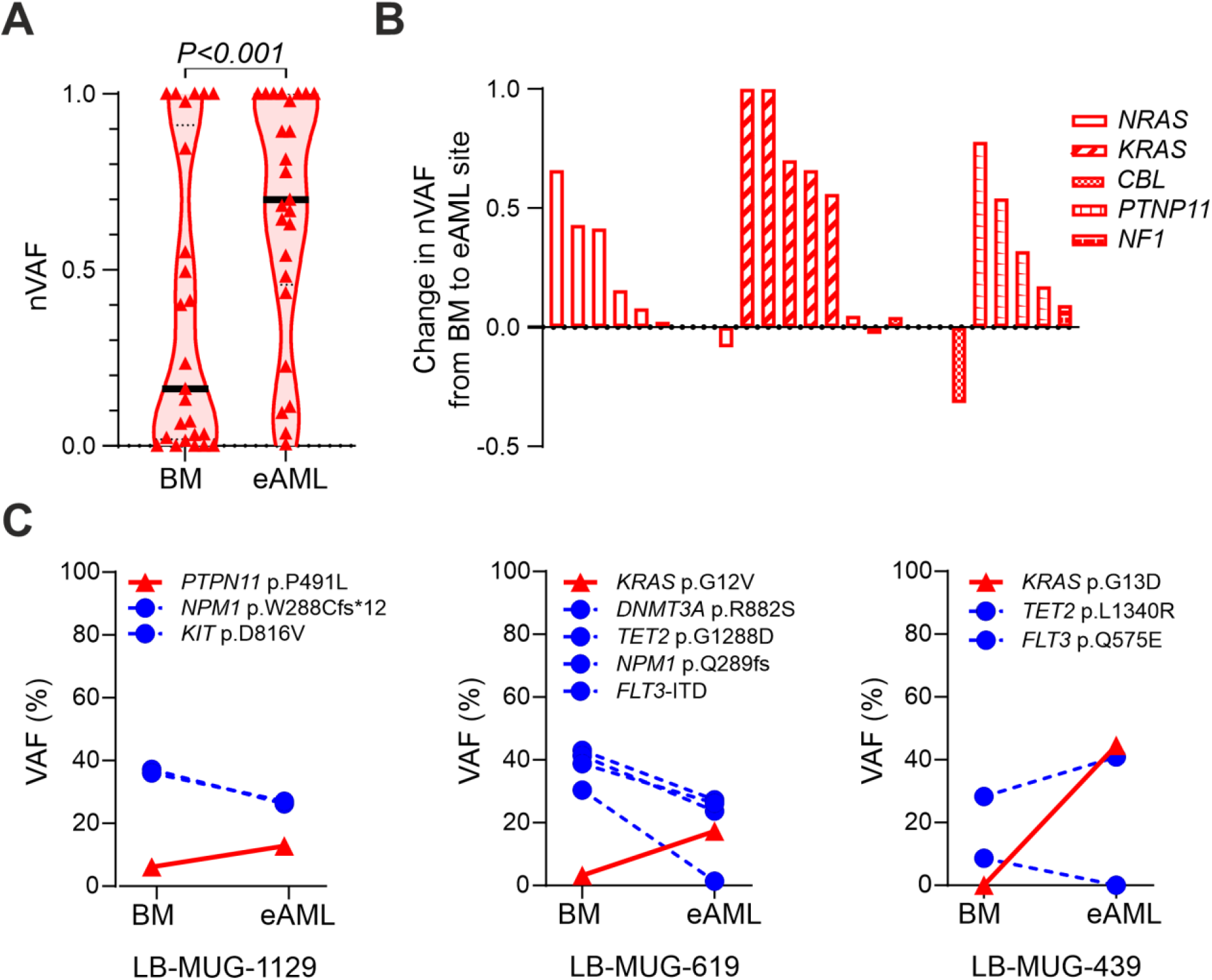
Clones with *RAS* and *RAS*-modifying mutations *(RAS*^*MUT*^) expand at extramedullary sites. (A) Violin plot of normalized variant allele frequencies (nVAFs) in 25 eAML cases for which paired bone marrow (BM) samples and eAML tissue biopsies were available for next-generation sequencing. nVAFs (VAF of a given *RAS*^*MUT*^ normalized to the highest VAF mutation within the same sample) were calculated to correct for differences in tumor purity between BM and eAML specimens^32^. Black horizontal lines indicate the median nVAF. Differences between groups were analyzed using the Mann–Whitney U test. (B) Waterfall plot illustrating the changes in nVAF between BM and eAML (nVAF_eAML_ – nVAF_BM_) across the 25 paired specimens. Positive values indicate enrichment of the *RAS*^*MUT*^ at the eAML site whereas negative values indicate higher VAFs in BM. (C) Representative individual paired eAML cases showing changes in *RAS*^*MUT*^ VAF between BM and eAML in the context of co-occurring mutations. Note that VAF values without normalization are shown in these cases. *RAS*^*MUT*^ are highlighted in red and marked with triangles (▲), whereas non-*RAS*^*MUT*^ are shown in blue and marked with circles (•). VAF, variant allele frequency; nVAF, normalized variant allele frequency; BM, bone marrow; eAML, extramedullary acute myeloid leukemia; LB-MUG, Leukemia Biobank-Medical University of Graz (Unique Biobank patient identifier).

### *RAS*^*MUT*^ drive the tissue infiltration of myeloid blasts and extramedullary leukemia growth

Given the enrichment of *RAS*^*MUT*^ in eAML biopsies, we next aimed to clarify the functional role of *RAS*^*MUT*^ in the tissue infiltration of myeloid blasts and extramedullary leukemia growth. Therefore, we introduced the *NRAS*^*G12D*^ mutation in the *RAS*-wildtype (WT) human myeloid leukemia cell line K562 to establish an isogenic cell line model. This mutation was chosen, as it was the most frequently detected alteration in the primary eAML specimens (Figure 1B). We introduced the *NRAS*^*G12D*^ mutation at the endogenous gene locus by adopting our previously established, highly active knock-in (KI) strategy using CRISPR/Cas9 and recombinant adeno-associated virus serotype 6 (rAAV6; Figure 3A; Supplemental Figure 1A).^18,33^ Importantly, a spleen focus-forming virus (SFFV)-driven fluorescent reporter expression cassette was integrated downstream of the cDNA to enable purification and tracking of modified cells via flow cytometry (GFP for *NRAS*^*G12D*^, mCherry and BFP for *NRAS*^*WT*^). The *NRAS*^*G12D/WT*^ fraction consisted of cells double positive for GFP and BFP, ensuring the presence of both wildtype and mutated alleles. For the *NRAS*^*WT/WT*^ population, cells double-positive for mCherry and BFP were sorted. In addition, we confirmed site-specific and seamless integration of *NRAS* cDNA, as well as the presence or absence of *NRAS*^*G12D*^ by In-out-PCR and sequencing respectively (Supplemental Figure 1B-C). Total NRAS protein levels were not altered, suggesting an intact cell-intrinsic regulation of gene expression (Supplemental Figure 2A). Further characterization of the cells revealed that *NRAS*^*G12D*^ activated the PI3K/AKT pathway, whereas MAPK/ERK-signaling remained largely unchanged (Figure 7A and Supplemental Figure 2B). The percentage of cells in S-Phase did not substantially differ between *NRAS*^*G12D/WT*^ and *NRAS*^*WT/WT*^ (Supplemental Figure 2C), validating previous observations that oncogenic *NRAS* in hematopoietic cells primarily affects cellular fitness and self-renewal rather than directly increasing proliferation.^34,35^

**Figure 3.**
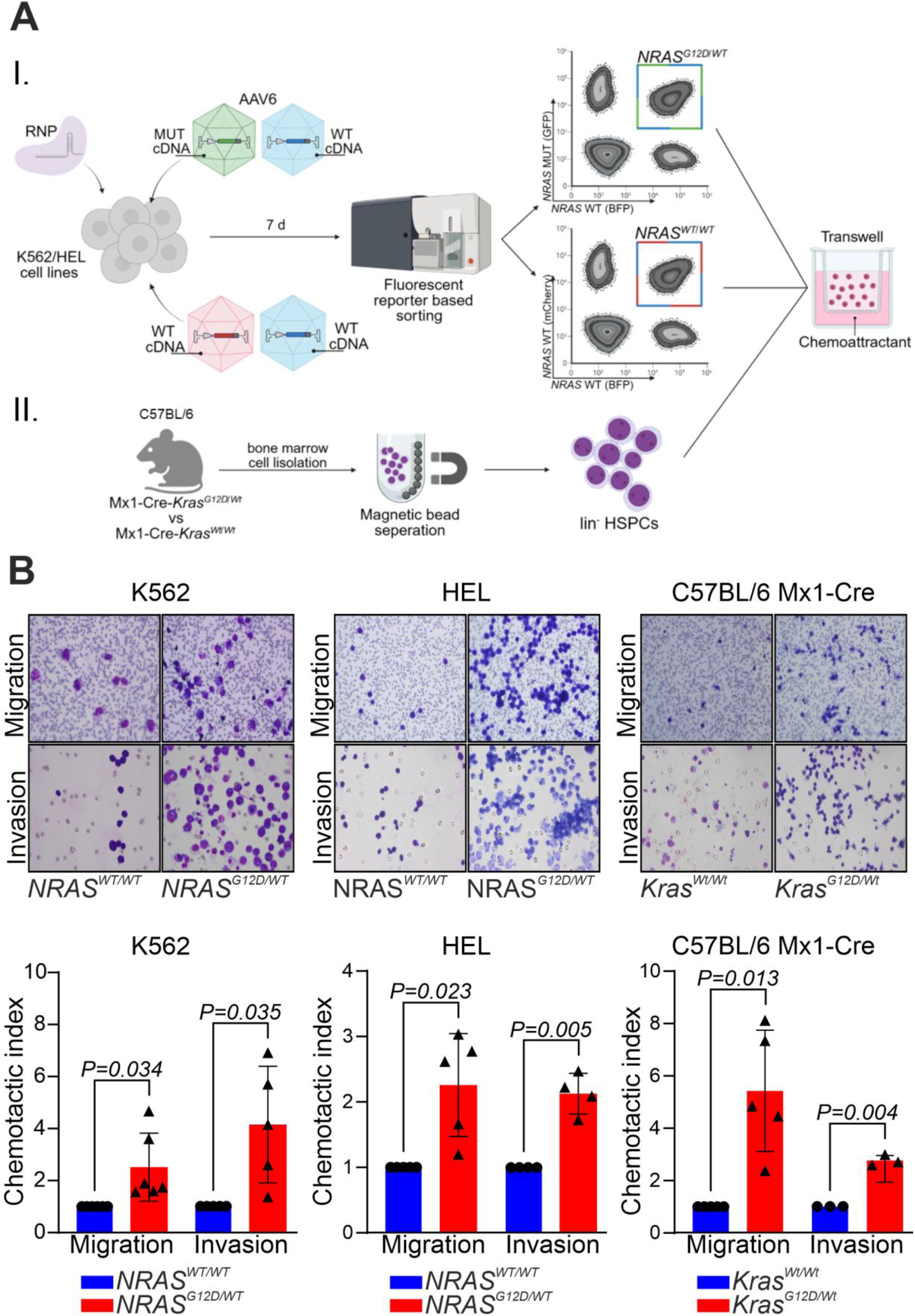
*RAS* and *RAS*-modifying mutations *(RAS*^*MUT*^) increase migration and invasion of leukemic cells in vitro. (A) Schematic overview of the generation of isogenic *RAS*^*MUT*^ and *RAS*^*WT*^ human and murine leukemic models and the subsequent in vitro transwell migration and invasion assays. (I) The hotspot *NRAS*^*G12D*^ mutation was introduced into K562 and HEL cell lines by CRISPR-Cas9 knock-in editing. Three donor AAV6 viruses were generated to create heterozygous genotypes. *NRAS*^*G12D*^ cDNA was linked to GFP and *NRAS*^*WT*^ cDNA was linked to BFP or mCherry. The *NRAS*^*G12D/WT*^ genotype was validated by double-positive GFP and BFP expression, whereas *NRAS*^*WT/WT*^ cells expressed BFP and mCherry. Correct bi-allelic fractions were isolated by fluorescence-based sorting. (II) For the murine model, bone marrow was harvested from *Mx1-Cre-Kras*^*G12D/Wt*^ and *Mx1-Cre-Kras*^*Wt/Wt*^ mice on C57BL/6 background, followed by lineage depletion to obtain lineage negative (lin^-^) hematopoietic stem and progenitor cells (HSPCs). (B) Transwell migration and invasion assays showing representative images (100× magnification, upper panels) and quantitative analysis (lower panels) of cells migrating through the membrane. In bar graphs (mean of all experiments +/- standard deviation), *RAS*^*WT*^ conditions (highlighted in blue and marked with circles for individual experiments [•]) were used as a calibrator and set to a value of 1. *RAS*^*MUT*^ conditions (highlighted in red and marked with triangles for individual experiments [▴]) are displayed as x-fold change relative to the calibrator. Differences between the groups were calculated using the one-sample *t*-test. CRISPR, clustered regularly interspaced short palindromic repeats; RNP, ribonucleoprotein; AAV6, adeno-associated virus serotype 6; MUT, mutant; WT, wildtype; cDNA, complementary DNA; mCherry, monomeric cherry fluorescent protein; GFP, green fluorescent protein; BFP, blue fluorescent protein; d, days; lin^-^ HSPCs, lineage negative hematopoietic stem and progenitor cells.

We then assessed the potential of these cells to migrate and invade in vitro. K562-*NRAS*^*G12D/WT*^ showed significantly higher rates of migration (P=0.034) and invasion (P=0.035) compared to K562-*NRAS*^*WT/WT*^ cells (Figure 3B). To exclude cell line specific effects, we validated these results in the human erythroleukemia cell line HEL. Again, HEL-*NRAS*^*G12D/WT*^ had a higher rate of migration (P=0.023) and invasion (P=0.005) compared to HEL-*NRAS*^*WT/WT*^ (Figure 3B). Moreover, we isolated hematopoietic stem and progenitor cells from a murine transgenic myeloid leukemia model driven by *Kras*^*G12D*^ (*Mx1-Cre-Kras*^*G12D/Wt*^) and respective control mice (*Mx1-Cre-Kras*^*Wt/Wt*^) and compared the cells in migration and invasion assays (Figure 3B). In agreement with the *NRAS*^*G12D*^ data, *Mx1-Cre-Kras*^*G12D/Wt*^ HSPC had a significantly higher migration (P=0.013) and invasion rate (P=0.004) than HSPC derived from *Mx1-Cre-Kras*^*Wt/Wt*^ mice (Figure 3B).

Next, we sought to validate these findings in vivo and performed a chorioallantoic membrane (CAM) assay.^21,36^ In this model, we evaluated the potential of K562-*NRAS*^*G12D/WT*^ and K562-*NRAS*^*WT/WT*^ to invade the CAM of chicken embryos and form solid tumor masses within this membrane (Figure 4A). In agreement with our in vitro data, K562-*NRAS*^*G12D/WT*^ invaded significantly better and formed significantly larger tumors in the CAM than K562-*NRAS*^*WT/WT*^ (P=0.025; Figure 4B). Consistent with the in vitro cell cycle analysis of *NRAS*^*G12D*^-edited cells, staining for the Ki-67 proliferation marker did not differ between the infiltrating K562-*NRAS*^*G12D/WT*^ and K562-*NRAS*^*WT/WT*^ cells (Supplemental Figure 3). We then aimed to validate these data in a murine model by injecting the K562 cells subcutaneously into immunocompromised NRG nude mice. This way of administration was preferred over the intravenous route as we aimed to transfer the cells into an extramedullary environment (Figure 4C). Consistent with the CAM assay, K562-*NRAS*^*G12D/WT*^ exhibited better extramedullary growth than K562-*NRAS*^*WT/WT*^ (P=0.011), as assessed by ultrasound (Figure 4D). Of note, tumors did not spread to other extramedullary sites or the BM at the experimental endpoint, which was determined based on predefined ethical criteria.

**Figure 4.**
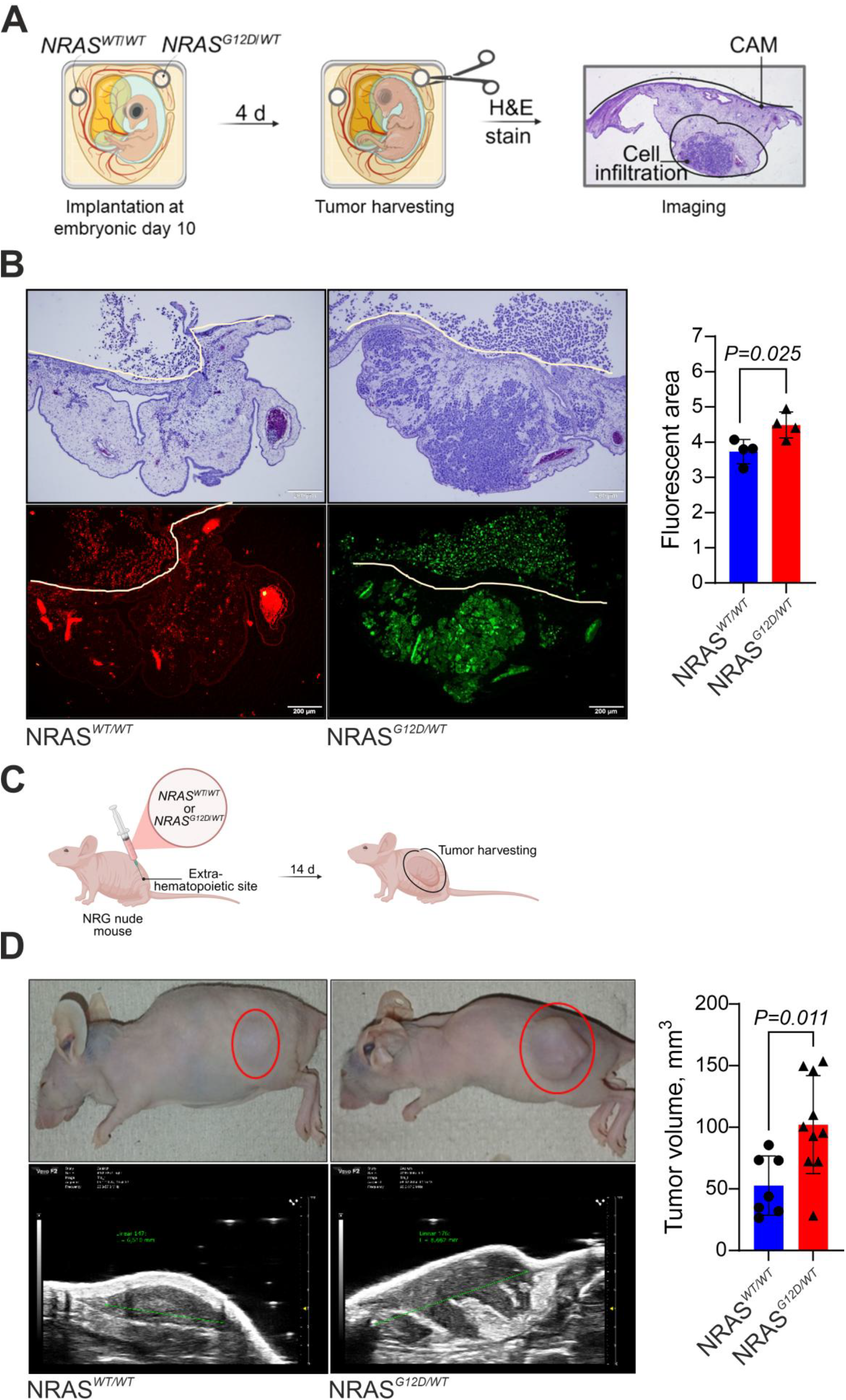
*RAS* and *RAS*-modifying mutations *(RAS*^*MUT*^) promote infiltration and the growth of leukemic cells at extramedullary sites. (A) Schematic overview of the chicken chorioallantoic membrane (CAM) assay. *RAS*-edited leukemic cells were seeded onto the CAM on day 10 of embryonic development and analyzed for CAM infiltration and tumor formation four days later. (B) Representative images of paraffin-embedded CAM sections following seeding of *NRAS*^*WT/WT*^ and *NRAS*^*G12D/WT*^ cells. Upper panels show hematoxylin and eosin (H&E) staining; lower panels show fluorescence (mCherry for *NRAS*^*WT/WT*^ and GFP for *NRAS*^*G12D/WT*^). The white line indicates the CAM membrane. The fluorescent area was quantified using ImageJ, background-corrected, and log10-transformed prior to statistical analysis. (C) Schematic workflow of extramedullary (subcutaneous) injection of *RAS*-edited leukemia cells into NRG nude mice. (D) Representative images of subcutaneous tumors (circled in red) at the time of sacrifice (upper panels) and corresponding ultrasound images (lower panels). The tumor volume (mm^3^) was assessed by ultrasound. In bar graphs (mean of all experiments +/- standard deviation), *NRAS*^*WT/WT*^ conditions are shown in blue and *NRAS*^*G12D/WT*^ in red. Individual experimental conditions are indicated by circles (•, *NRAS*^*WT/WT*^) and triangles (▴, for *NRAS*^*G12D/WT*^). Statistical differences between the groups were calculated using unpaired *t*-tests. d, days; WT, wildtype; NRG, NOD-*Rag1*^*null*^ *IL2rγ*^*null*^; tumor volume in mm^3^=[(length)x(width)x(height)]x(π/6).

### Tissue infiltration of *RAS*^*MUT*^ leukemic cells is driven through a *RAS*^*MUT*^-JAML-PI3K/AKT axis and is therapeutically amenable to AKT inhibition

To delineate the transcriptional changes associated with *RAS*^*MUT*^-driven leukemic tissue infiltration, we performed RNA-Seq of the subcutaneous *NRAS*^*G12D*^ and *NRAS*^*WT*^ tumors. This analysis revealed 258 differentially expressed genes with the majority (n=193) showing decreased expression in *NRAS*^*G12D/WT*^ tumors (Figure 5A). Gene set enrichment analysis showed that the downregulated genes were enriched for immune- and cytokine-related pathways, including interferon-γ response, inflammatory response, and allograft rejection (Supplemental Figure 4).

**Figure 5.**
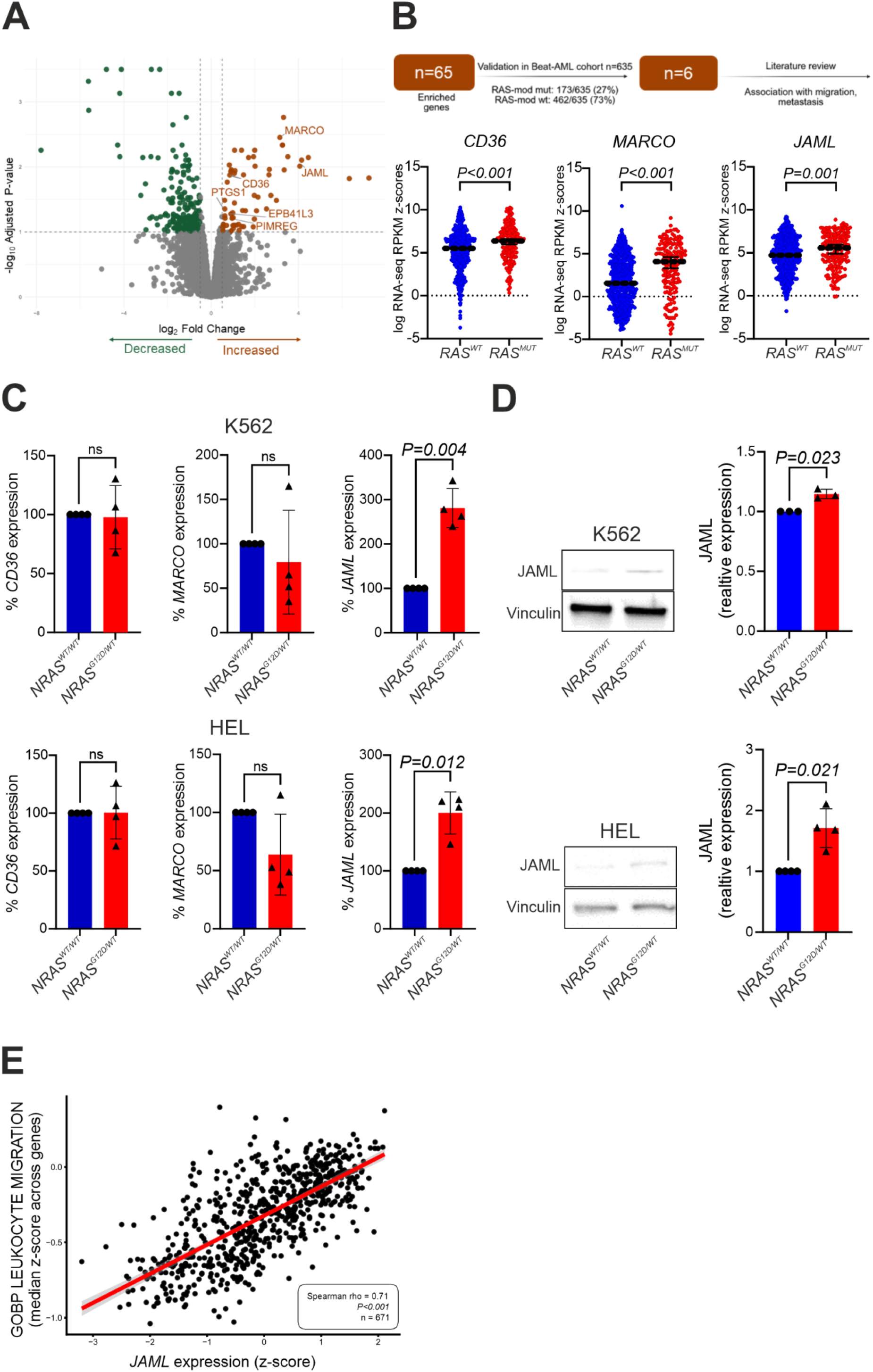
*JAML* expression is increased in leukemic cells with *RAS* and *RAS*-modifying mutations *(RAS*^*MUT*^). (A) Volcano plot depicting the differentially expressed genes in RNA Sequencing analysis in *NRAS*^*WT/WT*^ and *NRAS*^*G12D/WT*^ tumors derived from NRG nude mice. Each dot represents a different gene. Orange color indicates genes with increased expression in *NRAS*^*G12D/WT*^ tumors and green color indicate genes with decreased expression in *NRAS*^*G12D/WT*^ tumors. FDR (adjusted p-value) was set lower or equal to 0.1 and LFC higher or equal to +0.5 or lower or equal to -0.5. (B) Sixty-five genes with increased expression in *RAS*^*MUT*^ tumors were further evaluated in the Beat-AML cohort. Six of these genes (*CD36, EPB41L3, JAML, MARCO, PIMREG* and *PTGS1*) also showed increased expression in *RAS*^*MUT*^ AML patients within this cohort. Three candidates (*CD36, JAML*, and *MARCO*) have previously been implicated in cell migration and metastasis and were selected for further analysis. The graphs on the right show their expression z-scores in *RAS*^*WT*^ (blue) and *RAS*^*MUT*^ (red) patients. Medians are depicted by black horizontal lines, statistical differences between the groups were assessed by Mann Whitney tests. (C-D) Quantitative PCR (C) and Immunoblot analysis (D) showing selective upregulation of *JAML* across the *RAS*-modified cell line models. Gene expression was normalized to three independent reference genes (*GAPDH, B2M, PPIA*), and protein levels were normalized to Vinculin as a loading control. Values were subsequently normalized to the *RAS*^*WT*^ condition (set to 1). (E) Scatter plot showing the association between *JAML* expression and the median z score of genes annotated in the Gene Ontology Biological Process (GOBP) term “leukocyte migration”^26^ within the Beat-AML cohort.^24^ Each dot represents an individual sample (n=671). Red line indicates the fitted linear regression trend with 95% confidence interval (gray shading). A strong positive correlation was observed between *JAML* expression and leukocyte migration activity (Spearman ρ=0.71; P<0.001). In bar graphs (mean of all experiments +/- standard deviation) *RAS*^*WT*^ samples are shown in blue and marked with circles (•) and *RAS*^*MUT*^ samples in red and marked with triangles (▲), presented as fold change relative to the calibrator. Statistical comparisons were performed using one-sample *t*-tests. ns, non-significant; log, logarithm; MUT, mutant; WT, wildtype; JAML, junctional adhesion molecule-like protein; GOBP, gene ontology biological process; z-score, standard score; FDR, False Discovery Rate; LFC, Log2 Fold Change.

As we were mainly interested in identifying candidate genes amenable to direct therapeutic targeting, we focused on the 65 genes with increased expression in the *NRAS*^*G12D*^-mutated tumors. To refine this list, we analyzed their expression in the Beat-AML cohort,^24^ comparing patients with and without *RAS*^*MUT*^. Only six genes showed significantly higher expression in *RAS*^*MUT*^ cases (*CD36, EPB41L3, JAML, MARCO, PIMREG* and *PTGS1*). A literature search for invasion-, migration-, or metastasis-related genes highlighted *JAML*,^37,38^ *MARCO*,^39^ and *CD36*^40^ as potential candidates. Among these, only *JAML* showed consistently increased expression at both mRNA and protein levels in our experimental models (Figure 5C-D). JAML is a transmembrane adhesion protein, primarily involved in immune cell adhesion and migration, and facilitates leukocyte transmigration into tissue.^38^ Therefore, we examined the relationship between *JAML* expression and extramedullary disease. Notably, eAML status is not systematically annotated in the Beat-AML cohort. However, as extramedullary disease reflects leukemic cell dissemination, we assessed whether *JAML* expression is associated with transcriptional programs related to leukocyte trafficking. A leukocyte migration score was calculated for each sample as the median z-score of genes annotated to the Gene Ontology term GO:0050900 (leukocyte migration; n=671) and the values were correlated with *JAML* expression. These analyses revealed a strong positive correlation (Spearman rho=0.71; P<0.001), suggesting that *JAML* is linked to the activation of a dissemination-associated transcriptional program (Figure 5E). Based on these findings, we decided to further investigate the functional role of *JAML* in *RAS*^*MUT*^-driven tissue infiltration of leukemic blasts and established a *JAML* knockdown in K562-*NRAS*^*G12D/WT*^ by transfection of siRNA (Figure 6A-B). Indeed, the causative role of *JAML* expression could be corroborated by repeating the migration assays, where siRNA-mediated *JAML* knockdown in K562-*NRAS*^*G12D/WT*^ significantly decreased the migration potential of these cells (*JAML* siRNA1; P=0.016, *JAML* siRNA2; P=0.023; Figure 6C).

**Figure 6.**
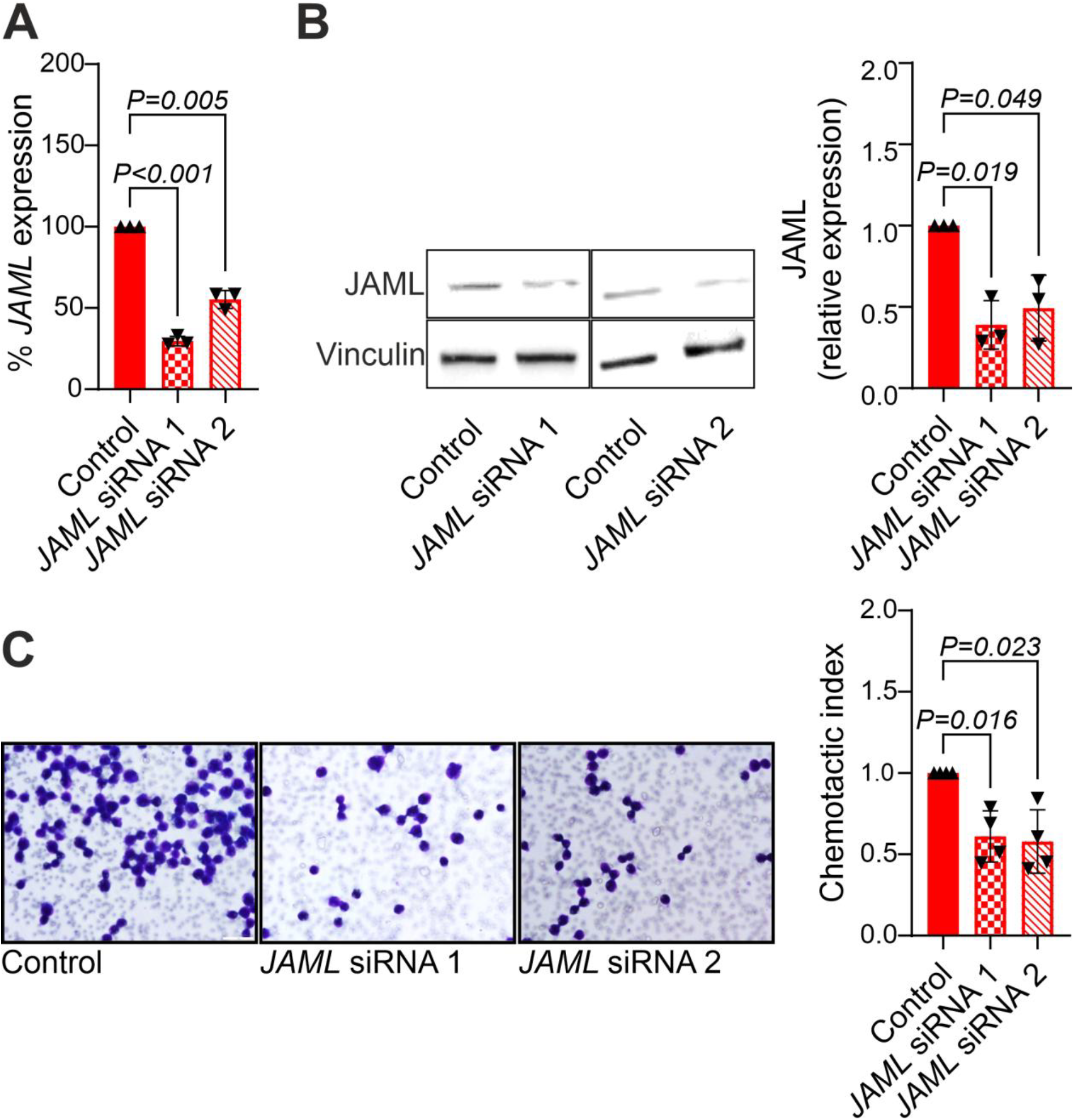
*JAML* knockdown reduces migration of leukemic cells with *RAS* and *RAS*-modifying mutations *(RAS*^*MUT*^). (A-B) siRNA-mediated knockdown of *JAML* in *NRAS*^*G12D/WT*^ K562 cells at the mRNA level (A, assessed by quantitative PCR) and protein level (B, assessed by Immunoblot). Gene expression was normalized to two independent reference genes (*GAPDH, B2M*), and protein levels were normalized to Vinculin as a loading control. Values were subsequently normalized to the control condition, which was transfected with non-targeting siRNAs and set to 1. (C) Transwell migration assays of K562-*NRAS*^*G12D/WT*^ cells following *JAML* knockdown. Representative images of Giemsa stained membranes were acquired at 100x magnification. In bar graphs, cells transfected with non-targeting control siRNA were used as the calibrator and set to a value of 1. *JAML* siRNA conditions are shown as x-fold change relative to the control. Bar graphs represent the mean of all experiments +/- standard deviation and symbols independent experiments (▴, control; ▾, *JAML* siRNA). Differences between the groups were calculated using the one-sample *t*-test. siRNA, small interfering RNA; JAML, junctional adhesion molecule-like protein.

Considering the *RAS*^*MUT*^-driven activation of the PI3K/AKT signaling cascade that we observed during the validation of successful CRISPR-Cas9-editing in K562/HEL-*NRAS*^*G12D/WT*^ cells (Figure 7A), we next investigated whether *JAML* expression is associated with PI3K/AKT signaling. Because phospho-protein data are not available in the Beat-AML cohort, we evaluated the relationship between *JAML* expression and a transcriptional signature consistent with PI3K/AKT pathway activity. A PI3K/AKT activation score was calculated for each sample as the median z-score of genes included in the HALLMARK_PI3K_AKT-MTOR_SIGNALING gene set (n=105) and correlated with *JAML* expression. *JAML* expression showed a strong positive correlation with this dataset (Spearman rho=0.43; P<0.001), consistent with a transcriptional program related to activated PI3K/AKT signaling (Supplemental Figure 5). Based on these findings, we next aimed to elucidate whether *JAML* indeed regulates this pathway. Two independent *JAML* siRNA constructs reduced AKT phosphorylation (Figure 7B) further supporting the existence of a *RAS*^*MUT*^-JAML-PI3K/AKT axis. The relevance of this signaling module in leukemic cell migration and tissue infiltration was further supported by migration assays: treatment of K562-*NRAS*^*G12D/WT*^ cells with the AKT inhibitor MK-2206 markedly decreased their migratory capacity (Figure 7C-D).

**Figure 7.**
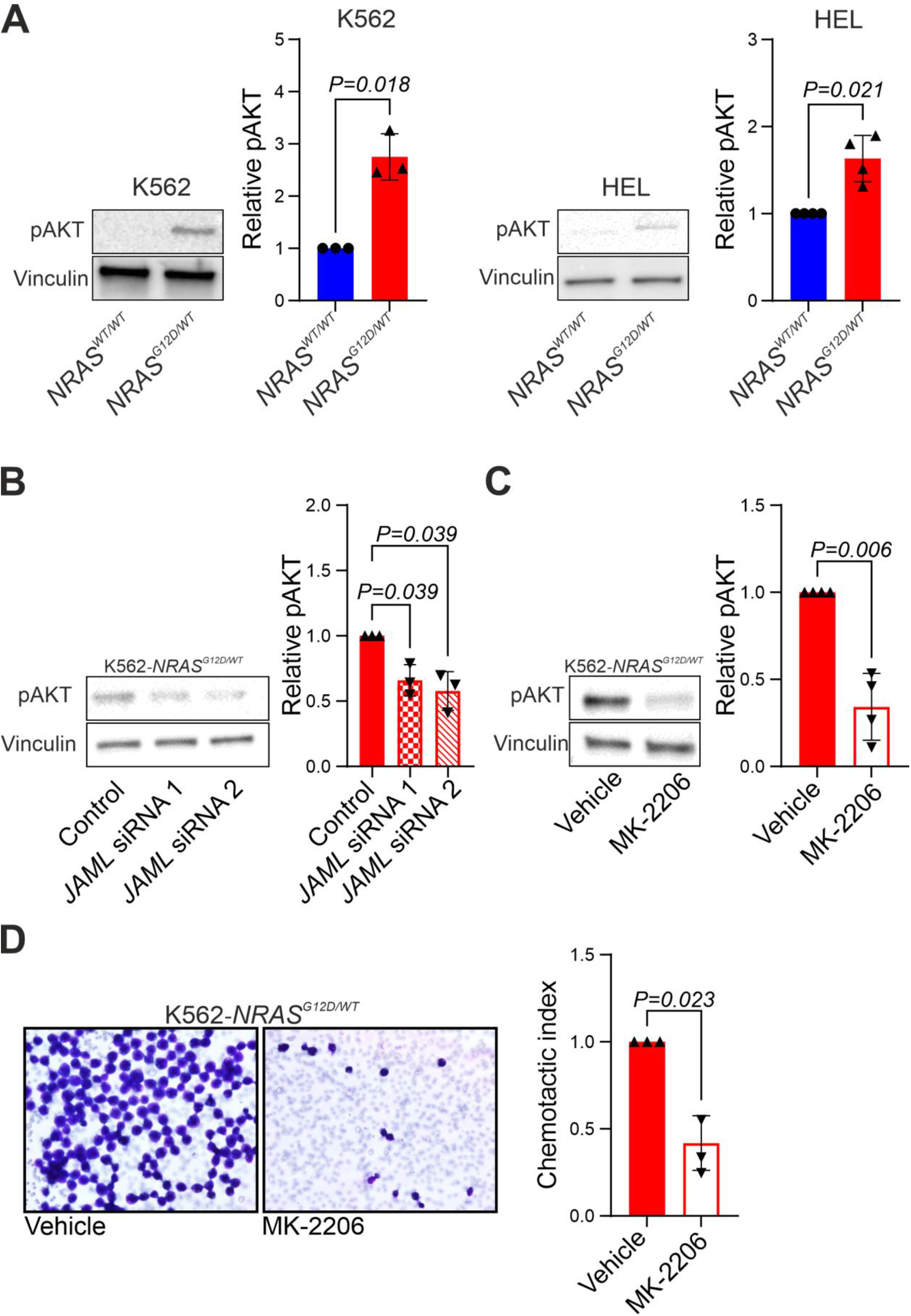
PI3K/AKT pathway inhibition reduces migration of *RAS* and *RAS*-modifying mutated *(RAS*^*MUT*^) cells. (A) Immunoblots depicting the activation of PI3K/AKT signaling pathway in CRISPR-edited in *NRAS*^*G12D/WT*^ cells. (B-C) Immunoblots showing decreased AKT phosphorylation as a surrogate marker for PI3K/AKT pathway inhibition in *NRAS*^*G12D/WT*^ cells following *JAML* knockdown (B) and pharmacological AKT inhibition with MK-2206 (C). pAKT levels were normalized to Vinculin as a loading control and subsequently normalized to control conditions (non-targeting siRNA in B; vehicle control [DMSO] in C). (D) Representative images of transwell migration assays acquired at 100x magnification. In bar graphs (mean of all experiments +/- standard deviation), *RAS*^*WT*^ samples are shown in blue and marked with circles (•) and *RAS*^*MUT*^ samples in red and marked with triangles (▴), presented as fold change relative to the calibrator. Symbols in the bar plots represent independent experiments. Control *RAS*^*MUT*^ conditions are indicated by triangles (▴) and *JAML* siRNA or MK-2206–treated samples by inverted triangles (▾). Statistical differences between the groups were calculated using the one-sample *t*-test. pAKT, phosphorylated AKT protein; WT, wildtype; JAML, junctional adhesion molecule-like protein; DMSO, Dimethyl Sulfoxide.

## Discussion

This study delineates the molecular landscape of 85 eAML cases, representing one of the largest genomic analyses of eAML to date. We show that *RAS*^*MUT*^ are enriched within this AML subform and affect more than 40% of eAML specimens. Analysis of paired eAML and BM specimens additionally revealed expansion or de-novo appearance of *RAS*^*MUT*^ clones at the extramedullary site, thereby highlighting potential limitations of BM-only molecular assessment in eAML patients. During the preparation of this manuscript, Nadorp et al. reported a multi-omic characterization of myeloid sarcoma and similarly identified activation of the *RAS* pathway as a recurrent feature of eAML.^41^ These findings are consistent with our observation that *RAS* pathway mutations are enriched in eAML and frequently expand at extramedullary sites. Together, these studies further support a central role of *RAS* pathway activation in the biology of eAML. To further elaborate on the role of *RAS*^*MUT*^ in tissue infiltration and eAML formation, we conducted functional studies employing in vitro migration and invasion assays and in vivo assays of eAML formation. Therefore, we introduced the most frequently detected *NRAS*^*G12D*^ mutation into *RAS*-WT myeloid leukemia cells, creating an isogenic cell line model. The CRISPR/Cas9-based strategy overcomes the bias of conventional transfection strategies, where the mutation is often expressed from an exogenous promoter and displays non-physiological overexpression.^42^ Using this approach, we show that *RAS*^*MUT*^ drives leukemic tissue infiltration and leukemic growth at extramedullary sites, strongly supporting the functional relevance of *RAS*^*MUT*^ in eAML development. These findings were validated using an additional human leukemia cell line and *Kras*-mutated HSPCs from a transgenic murine leukemia model, confirming their robustness in different cellular systems and species. They also place *RAS*^*MUT*^ in a small circle of other genetic aberrations with a proven functional role in eAML formation. Among others, these include *NPM1*^*43,44*^ and *DNMT3A*.^45^ Interestingly, *RAS*^*MUT*^ cells not only performed better than *RAS*^*WT*^ cells in invasion and migration in vitro but also increased the growth of leukemic cells at extramedullary sites in vivo. This finding is particularly remarkable, as we have shown that proliferation did not differ between the cells with and without *RAS*^*MUT*^, suggesting that other mechanisms underlie this phenotype. To obtain more mechanistic insights, we performed RNA-Seq and identified 258 differentially expressed genes between *RAS*^*MUT*^ and *RAS*^*WT*^ tumors, with the majority showing decreased expression in *RAS*^*MUT*^ tumors. Gene set enrichment analysis revealed a predominant downregulation of genes involved in immune regulation, including inflammatory responses, interferon gamma response, and allograft rejection. Similar changes have also been described in solid cancer models previously, suggesting that *RAS*-mutated leukemic cells may create an immunosuppressive environment in the extrahematopoietic niche, facilitating eAML growth.^46,47^ In this context, the system used may have limitations in fully addressing this question, as NRG nude mice are immunocompromised and lack T cells.^48^ However, they retain some innate immunity,^49^ which means immune dysregulation within the tumor microenvironment could still at least partly contribute to *RAS* mutation-driven eAML formation.

We then aimed to explore therapeutic implications. eAML can affect prognosis, and conventional therapeutic options often fail to clear eAML blasts.^3,4,6,7,10^ Particularly, *RAS*^*MUT*^ have been linked to resistance to targeted therapies in AML.^50,51^ For example, Wang et al. reported an AML patient with eAML skin infiltrates resistant to high-dose chemotherapy.^8^ BM and eAML sites showed discordant mutations, with *NRAS* mutation present only in the skin. Treatment with venetoclax/azacitidine (VEN/AZA) successfully controlled the disease in the BM, but leukemia cutis persisted, ultimately leading to *NRAS*^*MUT*^-driven systemic progression and death. Together, these findings and our data suggest that AML therapies should specifically target extramedullary clones to prevent eAML persistence and relapse. To identify potential therapeutic targets, we focused on the 65 genes with increased expression in *RAS*^*MUT*^ tumors. These genes were initially correlated to the *RAS*^*MUT*^ status within the Beat-AML cohort and only candidate genes with increased expression in *RAS*^*MUT*^ cases were further followed. Further validation via literature review, as well as validation by qPCR and immunoblot revealed the junctional adhesion molecule-like protein (JAML) as the most promising hit. More detailed analyses within the Beat-AML cohort revealed correlation of *JAML* expression with a transcriptional leukocyte migration program, which further strengthens its association to leukemic cell migration and tissue infiltration. Indeed, the causative role of *JAML* expression could be corroborated by repeating the migration assays, where siRNA-mediated *JAML* knockdown in K562-*NRAS*^*G12D/WT*^ significantly decreased the migration potential of these cells.

JAML is a transmembrane adhesion protein, primarily involved in immune cell adhesion and migration, and facilitates leukocyte transmigration into tissues.^38^ When expressed on cancer cells, it has been shown to increase cellular adhesion and interaction with the tumor microenvironment.^38^ Consistent with our study, *JAML* expression promotes tumor progression and metastasis in gastric cancer and lung adenocarcinoma.^37,52^ Unfortunately, specific targeting is not yet available for routine clinical use. It also remains unclear whether JAML inhibition would have to be used as a monotherapy or in addition to high-dose chemotherapy or non-intensive regimens, such as VEN/AZA. Furthermore, JAML has a dual-faceted role in tumorigenesis.^38^ Besides the above-described tumor-promoting effects on cancer cells, JAML expression on CD8+T cells mediates tumor-suppressing effects.^38^ In light of these data, establishing JAML as a therapeutic target in *RAS*^*MUT*^ eAML will require the development of an on-target delivery method that spares the normal tissues and particularly, the CD8+ T cell compartment, and is not foreseeable in the near future.

Of note, JAML has been described as activator of PI3K/AKT signaling.^37^ Although the exact mechanisms of JAML-driven AKT activation are unclear, JAML has been shown to directly interact with PI3K. Considering its transmembrane localization, JAML may help in translocating PI3K to the cell membrane, which is a well-established step in PI3K/AKT activation.^53,54^ As we observed activation of PI3K/AKT signaling in the *RAS*^*MUT*^ cellular models used in this study, we next analyzed the Beat-AML cohort and observed a correlation of *JAML* expression with transcriptional programs indicative of PI3K/AKT signaling. This is in line with our in vitro experiments, where *RAS*^*MUT*^ preferentially activated PI3K/AKT signaling, whereas MAPK/ERK signaling remained largely unchanged. This observation is in agreement with previous work from our group demonstrating that MEK inhibition does not inhibit the migration of *RAS*-mutated leukemia cells.^21^ Furthermore, recent multi-omic analyses of myeloid sarcoma failed to observe a correlation between *RAS* mutations and phospho-ERK.^41^ Together, these data suggest that *RAS* signaling in eAML may preferentially engage PI3K/AKT signaling rather than MAPK/ERK activation. Consequently, we investigated how *JAML* knockdown affects this pathway. Indeed, *JAML* siRNA-mediated silencing resulted in reduced AKT phosphorylation, supporting a *RAS*^*MUT*^-JAML-PI3K/AKT signaling axis involved in leukemic dissemination. Supporting this hypothesis, pharmacological inhibition of AKT with the orally available AKT inhibitor MK-2206 abrogated the *RAS*^*MUT*^-driven migration of leukemic blasts. Importantly, several AKT inhibitors (including MK-2206) have already been tested in clinical trials. Even more, the AKT inhibitor capivasertib was recently approved for treatment of breast cancer,^55^ demonstrating that pharmacologic AKT inhibition is clinically feasible. Therefore, our data highlight that AKT inhibition is a directly translatable therapeutic strategy for *RAS*^*MUT*^-driven eAML.

In conclusion, we have delineated the molecular landscape of eAML and show that *RAS*^*MUT*^ are associated with the development of this AML subform. We further show that *RAS*^*MUT*^ are functionally involved in the tissue infiltration of leukemic blasts and eAML formation. Finally, we show that upregulation of *JAML* and subsequent JAML-driven PI3K/AKT activation is an important contributor in *RAS*^*MUT*^-mediated eAML formation and represents an interesting therapeutic target for *RAS*^*MUT*^ eAML manifestations.

## Acknowledgments

This project was supported by Biobank Graz of the Medical University of Graz, Austria (Cohort 6002_14, Myeloid Neoplasm Collection). AZ is funded by the Austrian Science Fund (FWF, Grant-DOI 10.55776/P36672 and Grant-DOI 10.55776/PAT1753824), and the ERA-NET TRANSCAN-3 initiative (Austrian Science Funds Grant-DOI 10.55776/I6101). Research in his laboratory and Leukemia biobanking at the Medical University of Graz is further supported by Leukämiehilfe Steiermark. Students PC and AK are funded by the FWF within the PhD program Molecular Medicine of the Medical University of Graz. FS is funded by the ERA-NET TRANSCAN-3 initiative (BMFTR Grant No. 01KT2306B). AP is supported by the Austrian Science Fund (FWF, Grant-DOI 10.55776/P34109, Grant-DOI 10.55776/P29328, Grant-DOI 10.55776/FG30 and Grant-DOI 10.55776/PAT1140225) and by BioTechMed Graz. AR was supported by grants from the Austrian Science Fund (FWF, Grant-DOI 10.55776/P32783 and Grant-DOI 10.55776/I5021) and received support from the Austrian Society of Hematology and Oncology (Clinical Research Grant), Leukämiehilfe Steiermark and MEFOgraz. The Fig.s were in part created with Biorender (https://biorender.com). ChatGPT (OpenAI; https://chatgpt.com/) and Grammarly (https://www.grammarly.com/) were used to improve the clarity and grammar of the manuscript. All outputs were carefully reviewed, revised, and integrated by the authors, who take full responsibility for the final content.

## Authorship

Contribution: AZ designed and supervised the study; PC, JF, LH, BP, EG, AK, BB, KL, JF, KF, KG and KK performed the research; AZ, JF, SW, JN, SK, DW, GR, FS, AH, AW, HS and AR contributed patient specimens and collected clinical data; PC, JF, EG, JF, NGTW, BR, KF, KG, GH, KVC, GP, KK and AP collected data and performed statistical analyses; PC, JF, BP, EG, AK, JF, BR, KF, KG, NGTW, GH, KK, AP; FS, AH, AW, HS, AR and AZ analyzed and interpreted the data; PC and AZ wrote the manuscript; All authors read, revised and approved the manuscript.

