## Supplemental figures, tables and methods for "*RAS*-mutant clones drive extramedullary acute myeloid leukemia"

Chaida P, et al.

**Supplemental Figures and Tables**

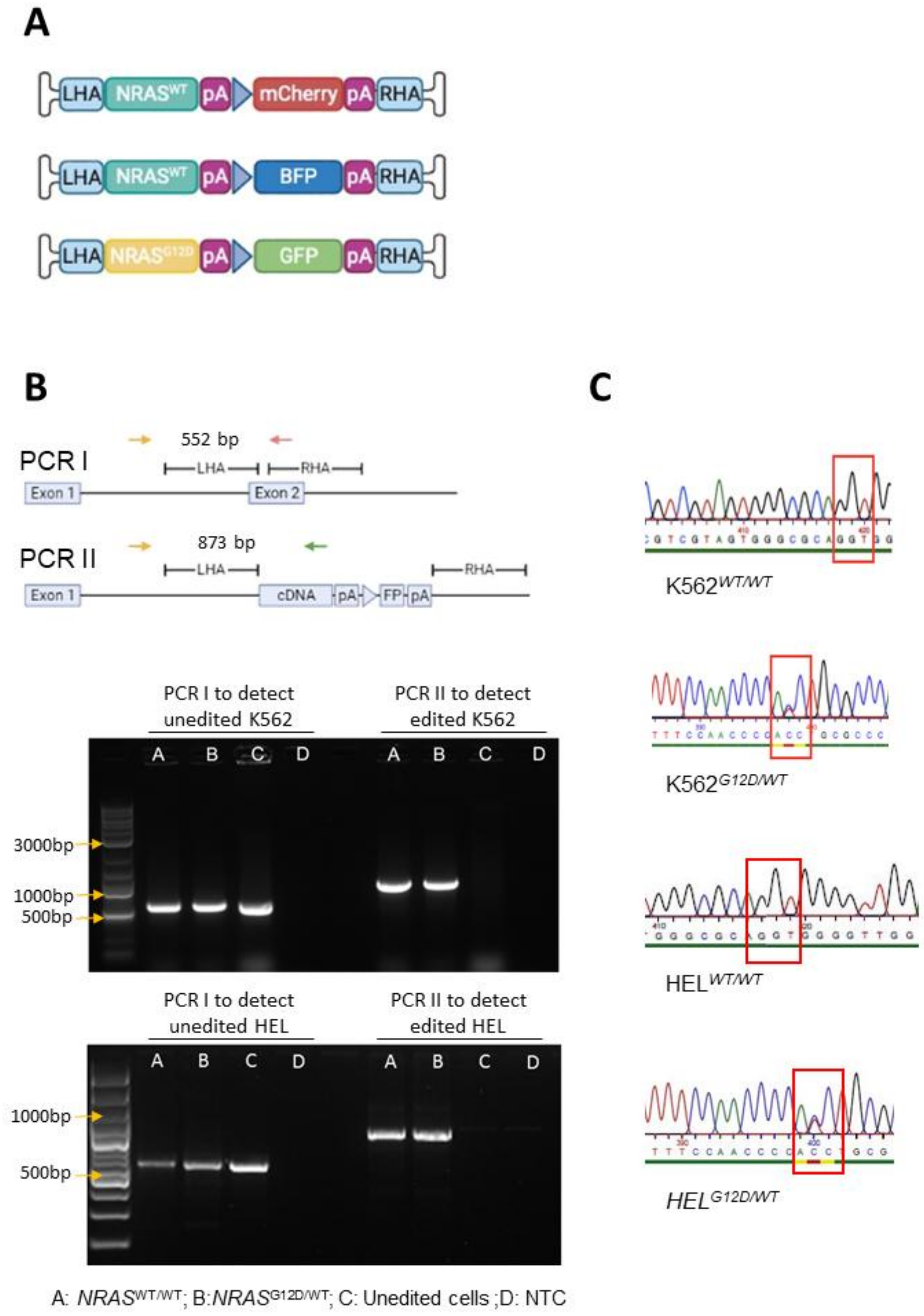


**Supplemental Figure 1. CRISPR/Cas9 editing cassette and validation on genomic level.** (A) Schematic representation of the donor template containing the whole *NRAS*$^{G12D}$ or *NRAS*$^{WT}$ codon optimized cDNA, followed by a simian-virus-40 polyadenylation polyadenylation (pA) signal, a cytomegalovirus (CMV) promoter, the fluorescent protein encoding gene, and a bovine growth hormone polyadenylation (pA) signal. In every case the donor template is flanked by the left homology arm (LHA) and right homology arm (RHA) both 400 bp. The three different donor templates were created to achieve heterozygous *NRAS*$^{WT/WT}$ and *NRAS*$^{G12D/WT}$ genotypes. *NRAS*$^{WT}$ cDNA was combined with either mCherry (monomeric Cherry) or BFP (blue fluorescent protein), while the *NRAS*$^{G12D}$ cDNA with GFP (green fluorescent protein). (B) In-out PCR schematic overview to detect the donor integration (upper panel). To detect unedited cells, the forward primer is located in the intron, right before the LHA and the reverse primer binds inside the RHA, resulting in a 552bp product (PCR I). The forward and reverse primers are depicted as yellow and red arrows respectively. For the detection of edited cells, the PCR utilizes the same forward primer as for the unedited cells. The reverse primer, however, binds inside the optimized cDNA, which is only present in successfully edited cells to avoid off-target binding. This PCR reaction yields an 873bp amplicon (PCR II). The primers are depicted as yellow (forward) and green (reverse) arrows. At the lower panel the agarose gel electrophoresis show the In-out-PCR products detecting edited or unedited alleles in K562 and HEL cells. On the gel position “A” *NRAS*$^{WT/WT}$ cells were loaded, on position “B” *NRAS*$^{G12D/WT}$ cells, position “C” has parental unedited cells and position “D” has the non-template control. (C) Sanger sequencing chromatograms confirming introduction of the G12D point mutation in K562 and HEL cells. The red box highlights the *NRAS*$^{G12D}$ locus. For the *NRAS*$^{WT/WT}$ genotype, a distinct peak within the G12 codon was visible, while for the *NRAS*$^{G12D/WT}$ a mixed signal occurred within the same locus. The latter confirms the correct introduction of the two gene cassettes, resulting in a cell line with at least one allele containing mutated *NRAS*$^{G12D}$ cDNA and at least one allele with *NRAS*$^{WT}$ cDNA. Green bars represent a quality >30%, yellow 20%-29%, red 10%-19% and black 0%-9%. Green peaks indicate Adenine, red Thymine, blue Cytosine and black Guanine. cDNA, complementary DNA; WT, wildtype; FP, fluorescent protein; PCR, polymerase chain reaction.

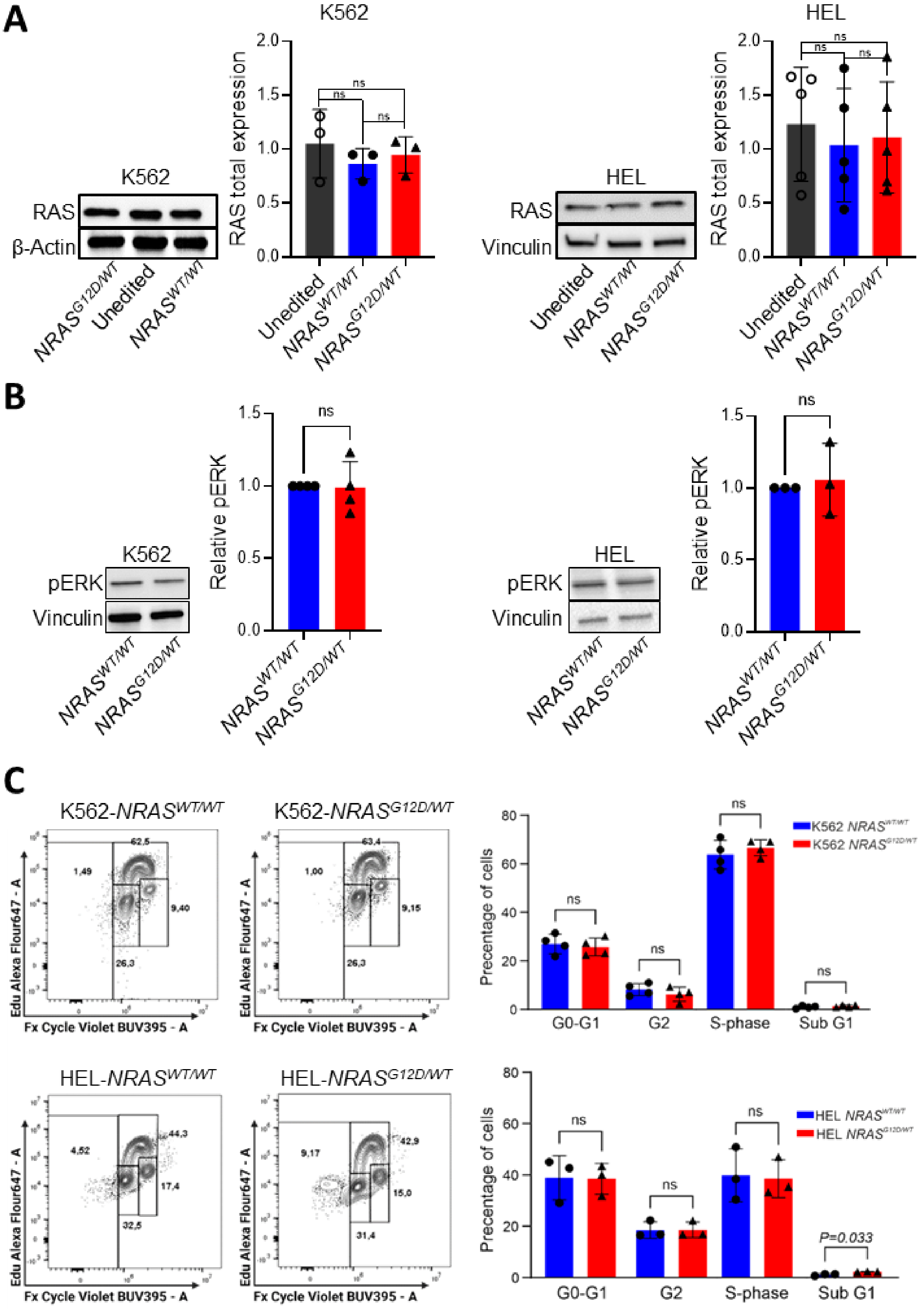


**Supplemental Figure 2. Functional validation of CRISPR/Cas9 editing in K562 and HEL cell lines.** (A) Immunoblots depicting the unaltered expression of total RAS protein between unedited and CRISPR-Cas9 edited *NRAS*$^{WT/WT}$ and *NRAS*$^{G12D/WT}$ cell lines, ensuring there was no introduction of stop codons as an off-target effect of editing. Total RAS protein levels were normalized to β-Actin or Vinculin as loading controls. (B) Immunoblots showing equal phosphorylation of ERK protein between *NRAS*$^{WT/WT}$ and *NRAS*$^{G12D/WT}$ cells. pERK levels were normalized to Vinculin as a loading control and subsequently normalized to control condition (*NRAS*$^{WT/WT}$ cells). (C) Flow cytometry panels and corresponding bar plots showing the percentage of the CRISPR-Cas9 edited K562 (upper panels) and HEL (lower panels) *NRAS*$^{WT/WT}$ and *NRAS*$^{G12D/WT}$ cells in each cell cycle phase (G0-G1; G2; S phase; Sub G1). In bar graphs (mean of all experiments +/- standard deviation) unedited cell lines are shown in grey and marked with white circles (○), *RAS*$^{WT}$ in blue and marked with circles (●) and *RAS*$^{MUT}$ samples in red and marked with triangles (▲). Statistical comparisons were performed using unpaired Student's t-tests and one-sample t-tests. WT, wildtype; pERK, phosphorylated ERK protein; ns, non-significant.

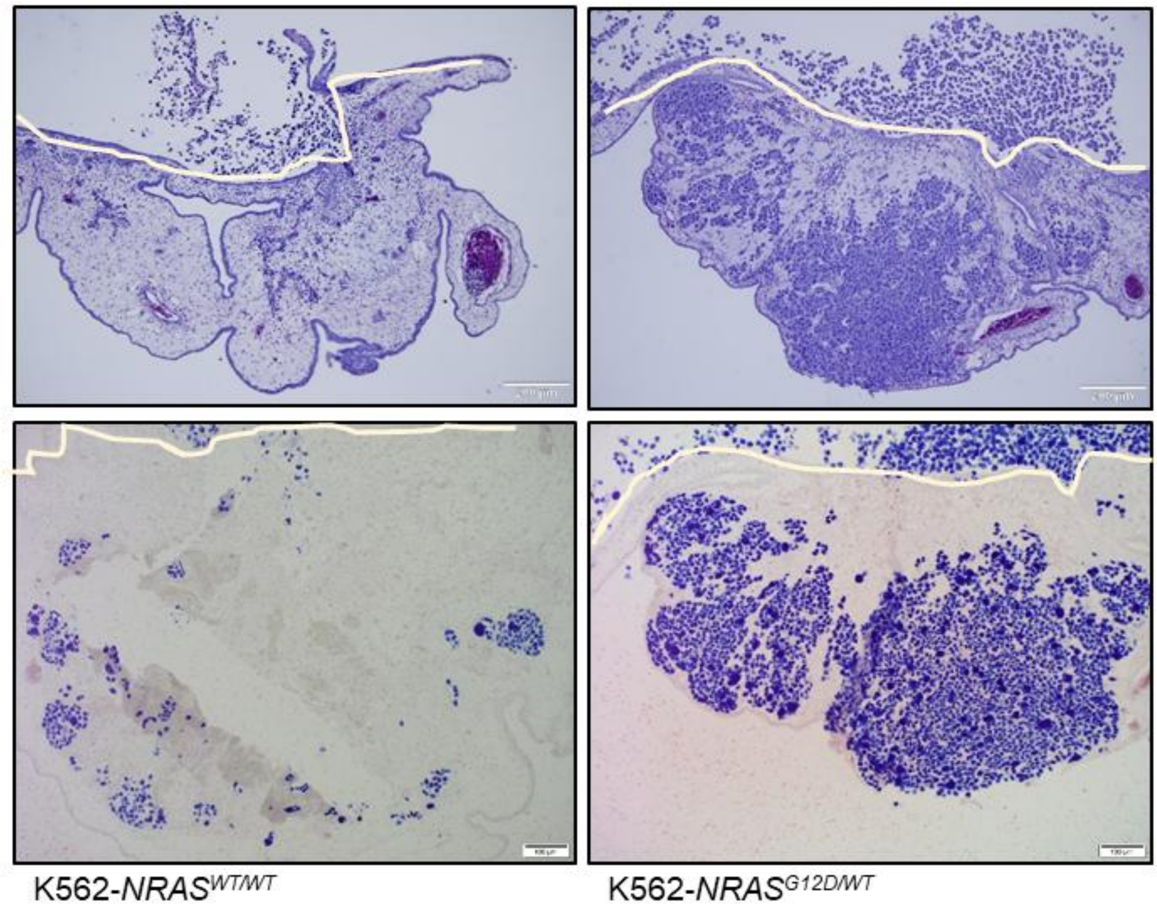


**Supplemental Figure 3. Proliferation assessment of the leukemic cells infiltrating the chorioallantoic membrane.** Tissue sections of K562-*NRAS*$^{WT/WT}$ and K562-*NRAS*$^{G12D/WT}$ tumors stained with hematoxylin & eosin (upper panel) and their corresponding immunohistochemistry staining with Ki67 proliferation marker (lower panel). The proliferating cells are stained with blue color. The white lines indicate the chicken chorioallantoic membrane border.

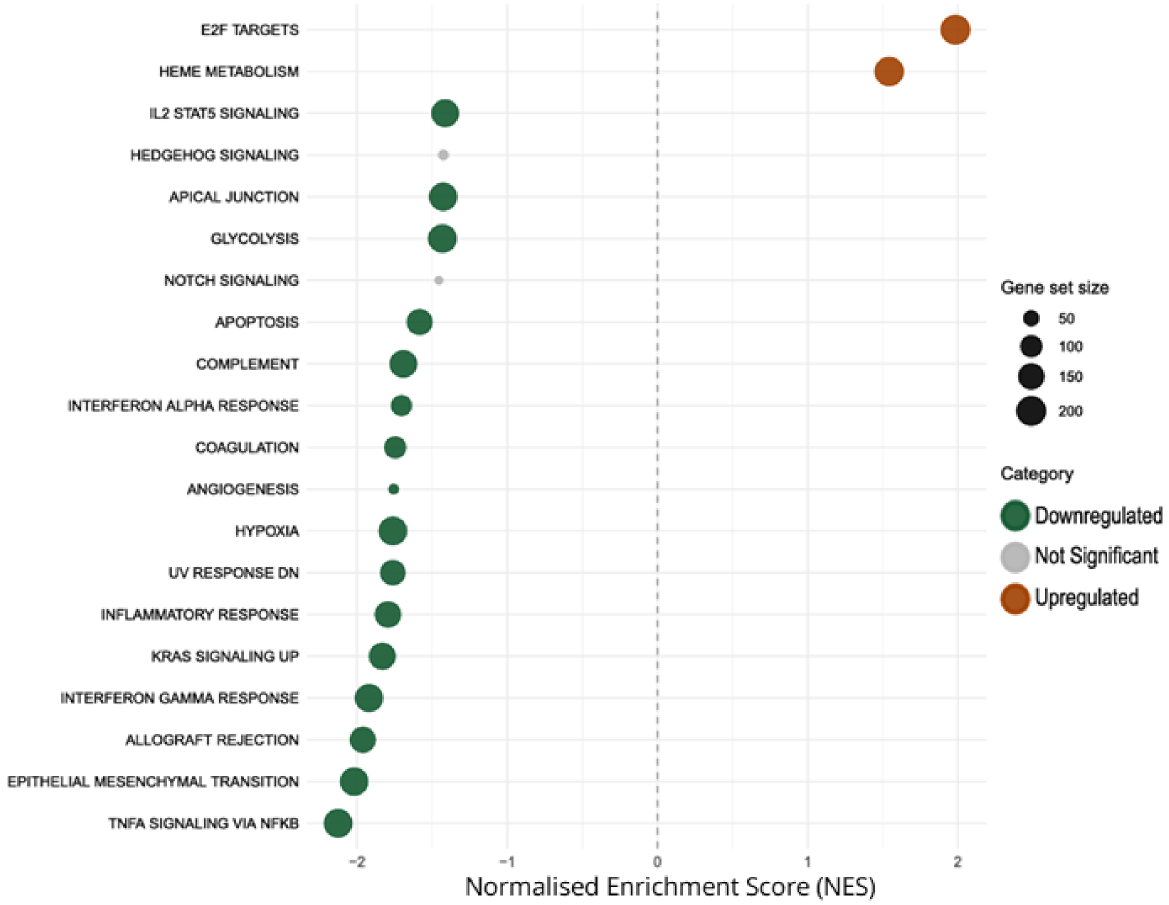


**Supplemental Figure 4. Gene set enrichment analysis (GSEA) identifies suppression of inflammatory and stress-response programs in *NRAS*$^{G12D/WT}$ tumors**. Bubble plot showing normalized enrichment scores (NES) for Hallmark gene sets ranked by differential expression between *NRAS*$^{WT/WT}$ and *NRAS*$^{G12D/WT}$ NRG-nude mice tumors. Positive NES (orange) indicates pathways enriched in *NRAS*$^{G12D/WT}$ tumors, whereas negative NES (green) indicates pathways enriched in *NRAS*$^{WT/WT}$ tumors. The most significantly enriched pathways are E2F targets and heme metabolism (as shown in orange bubbles). In contrast, multiple immune and inflammatory pathways demonstrate negative enrichment, including interferon-γ response, inflammatory response, and allograft rejection pathways (as shown in green bubbles). NES values were calculated using preranked GSEA, and significance was defined by false discovery rate (FDR) q value below the threshold. The order of the pathways was determined by NES score. Gray bubbles represent gene sets not meeting statistical significance thresholds.

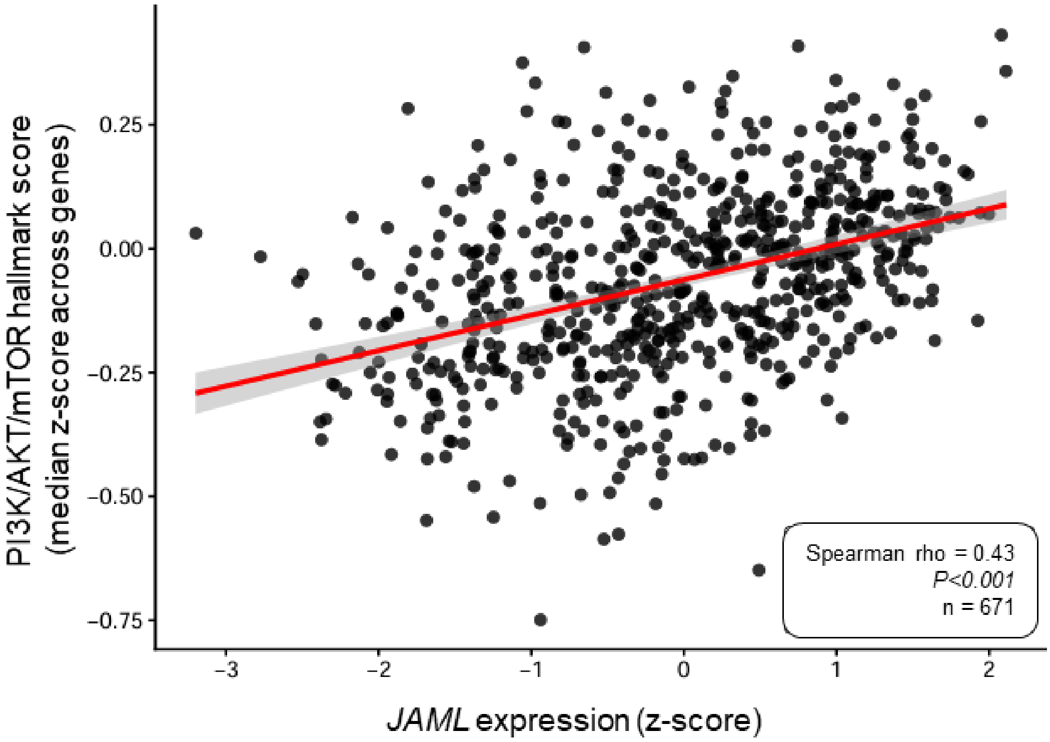


**Supplemental Figure 5. High *JAML* expression is correlated with a transcriptional signature consistent with PI3K/AKT pathway activity.** Scatter plot showing the correlation between *JAML* expression and the median z-score of genes included in the HALLMARK_PI3K_AKT-MTOR_SIGNALING gene set (n=105) within the Beat-AML cohort.[1] Each dot represents an individual sample (n=671). Red line indicates the fitted linear regression trend with 95% confidence interval (gray shading). *JAML* expression was positively correlated with pathway activity (Spearman ρ=0.43; P<0.001).

**Supplemental Table 1. Patient characteristics of the 85 eAML patients analyzed.**

| | |
|---|---|
| **Age in years, median (range)** | 63 (26-93) |
| **Sex, n=** | |
| Male | 37/85 (43.5%) |
| Female | 48/85 (56.5%) |
| **Isolated eAML, n=** | |
| Yes | 12/71 (16.9%) |
| No | 59/71 (83.1%) |
| Not available | 14 |
| eAML at relapse, n= | |
| Yes | 20/72 (27.8%) |
| No | 52/72 (72.2%) |
| Not available | 13 |
| **eAML site, n=** | |
| Skin | 51/85 (60.0%) |
| Lymph nodes | 11/85 (12.9%) |
| Bone | 4/85 (4.7%) |
| Soft tissue | 4/85 (4.7%) |
| Breast | 3/85 (3.5%) |
| Liver | 2/85 (2.4%) |
| Skeletal muscle | 2/85 (2.4%) |
| Pancreas | 2/85 (2.4%) |
| Brain | 1/85 (1.2%) |
| Stomach | 1/85 (1.2%) |
| Parotid gland | 1/85 (1.2%) |
| Oral mucosa | 1/85 (1.2%) |
| Pleura | 1/85 (1.2%) |
| Spleen | 1/85 (1.2%) |

Note that percentages may not total 100 due to rounding.

**Supplemental Table 2. *RAS*$^{MUT}$ status in paired bone marrow and eAML tissue samples**

| | |
|---|---|
| ***$RAS^{MUT}$*** **eAML specimens with corresponding bone marrow sequencing data available** | 25 |
| *$RAS^{MUT}$* new at eAML tissue, n= | 4/25 (16.0%) |
| ***$RAS^{WT}$*** **eAML specimens with corresponding bone marrow sequencing data available** | 40 |
| *$RAS^{MUT}$* lost at eAML tissue, n= | 0/40 (0%) |

### Supplemental Methods

#### Next generation sequencing (NGS) gene lists

NGS of the Austrian cohort comprised sequencing of the whole coding region of *BCOR, BCORL1, CEBPA, DDX41, DNMT3A, ELANE, ETNK1, ETV6, GATA2, GNB1, HAX1, NF1, PHF6, PIGA, PPM1D, PRPF8, SF3B2, SFRP1, SRP72, STAG2, TP53*, and *ZRSR2*, as well as mutational hotspot regions of *NPM1, ASXL1, BRAF, CALR, CBL, CSF3R, CXCR4, ETNK1, EZH2, FLT3, IDH1, IDH2, JAK2, KIT, KRAS, MPL, NRAS, PTPN11, RUNX1, SETBP1, SF3B1, SRSF2, STAT3, STAT5B, TET2, U2AF1* and *WT1* genes. NGS of the German cohort was performed using the Myeloid-NGS DNA Capture (Myeloid-NDC, Univ8 Genomics, Belfast, United Kingdom).

#### CRISPR/Cas9 mutational knock-in

The sgRNA targeting *NRAS* was designed to bind in the beginning of exon 2, with target sequence 5'-GACTGAGTACAAACTGGTGG -3'. The sgRNA was obtained from Synthego (Redwood City, CA, USA) with chemical modifications. The codon-optimized *NRAS* mutant or wild-type cDNA, spanning exons 2-5 and flanked by homology arms, was inserted into a pAAV-MCS plasmid (Agilent #240071, Santa Clara, CA, USA). Vectors also contained a fluorescent reporter gene (GFP, BFP, or mCherry) driven by an SFFV-promoter, positioned downstream of the *NRAS* cDNA. Recombinant AAV6 was produced as described previously.[2,3] K562 cells were electroporated using the Neon™ Transfection System (Thermo Fisher Scientific) and the Neon™ Transfection System 100µL Kit, (Thermo Fisher Scientific, Waltham, MA, USA), and HEL cells using the Lonza 4D Nucleofection system and the SF Cell line Nucleofection Kit L (Lonza, Cologne, Germany). Nucleofection conditions: $1\times10^6$ cells/ml, 15 µg Cas9 protein (IDT, Coralville, IA, USA) pre-complexed with sgRNA at 1:2.5 molar ratio, Lonza program FF-100. Following electroporation, the cells received 0.5 µM of AZD7648 inhibitor (MED Chem Express, Monmouth Junction, NJ, USA), to inhibit non-homologous end joining, thereby enhancing homology directed repair (HDR) mediated integration efficiency. The cells were then immediately transduced with 5000 vector genomes/cell of AAV6. Culture media was refreshed after 24h and cells exhibiting high reporter expression were sorted 7 days post-transduction usind a FACSAria IIIu (Beckman Coulter, San Jose, CA, USA). Correct donor template integration was confirmed by In-out PCR and Sanger sequencing across the junction and the edited target sites.

#### Characterization of editing

For characterization of editing on genomic level, In-out PCR was performed as previously described.[4]This technique employs three primers. The forward primer (5'-CTGGAGACAAAGGCCTTGGGCG-3') is the same for edited and unedited cells, and its binding site is located in the intron right before the left homology arm (LHA), but there are two different reverse primers. For the detection of unedited cells, one of the reverse primers (5'GGATCAGGTCAGCGGGCTACCA-3') is located within the right homology arm (RHA) resulting in a 552 bp PCR product. For the detection of edited cells, the second reverse primer (5'-GCGTCCTCTACACCCTGTCTAGT-3') is located within the gene cassette. This PCR reaction yields an 873 bp amplicon. To verify the correct heterozygous genotype, the product of the In-out-PCR for the detection of edited cells was sequenced. The chromatogram was examined for the sequencing quality of $NRAS^{WT/WT}$ and $NRAS^{G12D/WT}$.

Cell cycle analysis after editing was assessed using the Click-iT™ Plus EdU Flow Cytometry Assay Kit (Thermo Fisher Scientific, Waltham, MA, USA) according to the manufacturer's instructions. 5-ethynyl-2'-deoxyuridine (EdU), a thymidine analog, is incorporated into newly synthesized DNA during S phase and subsequently detected via a copper-catalyzed azide–alkyne cycloaddition reaction using an Alexa Fluor™ 647–conjugated azide. For DNA content analysis and cell cycle discrimination, FxCycle™ Violet stain was added. Analysis was performed on CytoFLEX LX flow cytometer (Beckman Coulter, San Jose, CA, USA). The percentage of cells in different cell cycle phases (G0/G1, S, G2/M) were determined based on FxCycle™ Violet DNA content in combination with EdU incorporation.

Downstream effects of $NRAS^{G12D}$ in MAPK/ERK and PI3K/AKT signaling pathways were evaluated with immunoblot for the detection of pERK and pAKT. The results were normalized to Vinculin. Densitometry was measured with ImageJ (https://imagej.net/ij/) software (Supplementary Figure 2C).

**Immunoblots and immunohistochemistry**

Immunoblot were performed as previously described[5–7] using the following antibodies: RAS, pERK, pAKT, JAML, β-Actin, Vinculin. ImageJ (https://imagej.net/ij/) software was used for densitometry. Immunhostochemistry was performed as described previously using the Ki67 antibody.[7]

**Mouse Hematopoietic Progenitor (Stem) Cell Enrichment**

Bone marrow cells were isolated from 8–9-week-old Mx1-Cre-$Kras^{G12D/Wt}$ mice exhibiting clinical features of myeloproliferative disease and age-matched C57BL6/J Mx1-Cre-$Kras^{Wt/Wt}$ siblings. Mice were euthanized, and femurs, tibias, hips, and spines were harvested. Bones were mechanically crushed using a mortar and pestle. Red blood cells were lysed using 1mL of 1×PharmLyse™ (BD Biosciences, San Jose, CA, USA) for 10 minutes at room temperature. Lineage depletion per $1x10^6$ cells was performed using "Mouse Hematopoietic Progenitor (Stem) Cell Enrichment Set - DM" (BD Biosciences), following the manufacturer's instructions. Lineage negative hematopoietic stem and progenitor cells were isolated through magnetic based separation as cells committed to the T- and B-lymphocytic, myeloid (monocytic and granulocytic), and erythroid lineages are depleted.

**RNA extraction, reverse transcription and real time quantitative PCR**

RNA was isolated from K562 and HEL edited hematopoietic cell lines employing the miRNAeasy Micro Kit (Qiagen, Hilden, Germany) following the manufacturer´s instruction. cDNA synthesis was performed with TaqMan Reverse Transcription (RT) Reagents (Applied Biosystems, Carlsbad, CA,USA) starting from 500 ng of total RNA and using random hexamers for RT. An Applied Biosystems 7500 Real-Time PCR System and the SYBR Green method (Invitrogen, Carlsbad, CA,USA) were used for real time quantitative PCR (qPCR) for *CD36* (Fwd primer 5'-ACGCTGAGGACAACACAGTC-3'; Rev primer 5'-GCCACAGCCAGATTGAGAAC-3'), *JAML* (Fwd primer 5'-CGCCTGAGCTAACAGTCCAT-3'; Rev primer 5'-

TCCTGGTGACAGAGTCCAGT-3’) and *MARCO* (Fwd primer 5’-TGGGGGACAATTTGCGATGA-3’; Rev primer 5’-CCACTTTGTACAGGGCCCTT-3’) expression analysis. DNA concentration was 2.5 ng/µl and *B2M, PPIA* and *GAPDH* were selected as control genes for the normalization and analysis performed using the ΔΔCt method as described previously.[5,8,9]

**RNA extraction, library preparation and NovaSeq Sequencing**

Total RNA was isolated from frozen tumor tissues of K562 $NRAS^{WT/WT}$ and $NRAS^{G12D/WT}$ cells, extracted from NRG-nude mice, using Qiagen RNeasy Mini kit following manufacturer’s instructions (Qiagen, Hilden, Germany). RNA samples were quantified using Qubit 4.0 Fluorometer (Life Technologies, Carlsbad, CA, USA) and RNA integrity was checked with RNA Kit on Agilent 5300 Fragment Analyzer (Agilent Technologies, Palo Alto, CA, USA). RNA sequencing libraries were prepared using the NEBNext Ultra II RNA Library Prep Kit for Illumina following manufacturer’s instructions (NEB, Ipswich, MA, USA), using 100ng RNA per sample as input. Sequencing libraries were validated using NGS Kit on the Agilent 5300 Fragment Analyzer (Agilent Technologies, Palo Alto, CA, USA), and quantified by using Qubit 4.0 Fluorometer (Invitrogen, Carlsbad, CA). The sequencing libraries were multiplexed and loaded on the flowcell on the Illumina NovaSeq X plus instrument according to manufacturer’s instructions. The samples were sequenced using a 2x150 Pair-End (PE) configuration v1.5. Image analysis and base calling were conducted by the NovaSeq Control Software v1.7 on the NovaSeq instrument. Raw sequence data (.bcl files) generated from Illumina NovaSeq was converted into fastq files and de-multiplexed using Illumina bcl2fastq program version 2.20. One mismatch was allowed for index sequence identification.

**Gene set enrichment analysis**

Differential gene expression and Gene Set Enrichment Analysis (GSEA) analyses were performed using a RNA-SAIL (v.2.0.0) R package, available on GitHub at (https://github.com/OncologyMedunigraz/rna-sail) with default settings LFC threshold higher or equal to +0.5 or lower or equal to -0.5, and FDR of 0.1.

**siRNA transfection**

Transient downregulation of *JAML* (*AMICA-1*) was performed in K562-$NRAS^{G12D/WT}$ cells with two different ON-TARGETplus siRNA sequences Dharmacon™ (Dharmacon, Lafayette, CO; Revvity, Waltham, MA, USA) at 150nM, along with a non targeting siRNA as a negative control. Target sequences: siRNA-1 5’-GACCAAGGUAGAAUGGAUA-3’, siRNA-2 5’-ACGAAUAUGUGCUAUACUA-3’. siRNA was introduced into the cells via electroporation (Lonza 4D Nucleofection system) suspended in 4D Nucleofector X Optimization Kit for Cell Lines (Lonza, Cologne, Germany), program FF-120. Cells were then cultured with RPMI-1640 enriched with 10% FBS without antibiotics. Expression of JAML was analysed by qPCR and immunoblot 24h-96h post transfection. RNA extraction, reverse transcription and qPCR were performed as described above using *B2M* and *GAPDH* for normalization. Immunoblot validation for JAML downregution was performed using Vinculin for normalization and ImageJ (https://imagej.net/ij/) software for densitometry analysis.
